# Hi-D: Nanoscale mapping of nuclear dynamics in single living cells

**DOI:** 10.1101/405969

**Authors:** Haitham A. Shaban, Roman Barth, Ludmila Recoules, Kerstin Bystricky

**Affiliations:** Laboratoire de Biologie Moléculaire Eucaryote (LBME), Centre de Biologie Intégrative (CBI), CNRS; University of Toulouse, UPS; 31062 Toulouse; France; Spectroscopy Department, Physics Division, National Research Centre, Dokki, Cairo, Egypt

## Abstract

Bulk chromatin motion has not been analysed at high resolution. We present Hi-D, a method to quantitatively map dynamics of chromatin and abundant nuclear proteins for every pixel simultaneously over the entire nucleus from fluorescence image series. Hi-D combines reconstruction of chromatin motion, and classification of local diffusion processes by Bayesian inference. We show that DNA dynamics in the nuclear interior are spatially partitioned into 0.3 – 3 μm domains in a mosaic-like pattern, uncoupled from chromatin compaction. This pattern was remodelled in response to transcriptional activity. Hi-D can be applied to any dense and bulk structures opening new perspectives towards understanding motion of nuclear molecules.

## INTRODUCTION

Spatial organization and dynamics of chromatin correlate with cell function and fate [1]. On the coarsest level, chromosomes occupy territories in mammalian cells [2]. The relative proportion of dense heterochromatin and open euchromatin regions reflect cellular activity [3]. Transitions within and between eu- and heterochromatin involve the remodeling of multiple hierarchical levels of chromatin organization from domain folding and long-range looping to nucleosome density to adapt to and enable DNA processing [4]. Structural models derived from contact and crosslinking frequencies [5–7] are consistent with the view that the genome is partitioned into functional compartments and sub-compartments [8]. It is now becoming increasingly clear that such nuclear compartments are also dynamic entities whose conformational changes impact mechanisms and function of genome folding [9]. Tracking of labelled single DNA loci [10–14] or chromatin domains [15,16] demonstrated that chromatin motion is highly heterogeneous at short time intervals. Sparse loci, however, are difficult to place in the context of global chromatin organization [17] and locally restrained genomic processes can also not be inferred from quantities averaged over the entire nucleus [18,19]. A first study analyzing bulk chromatin motion at nanoscale resolution revealed dynamic partitioning of chromatin into a number of nuclear sub-regions with correlated motion of chromatin in the micrometer range [20]. However, the cause and/or effect of (correlated) chromatin dynamics is not yet clear. Likewise, whether compaction of chromatin determines its spatial coherence or whether chromatin dynamics are distinct in open and closed chromatin is still a matter of debate.

To tackle this need, we developed a new approach called High-resolution Diffusion mapping (Hi-D) that overcomes the limitations of sparse and ensemble approaches. Hi-D combines a dense Optical Flow reconstruction to first quantify the local motion of chromatin and other abundant nuclear constituents at sub-pixel accuracy within a series of images, and a Bayesian inference approach in the second step to precisely classify local types of diffusion. Biophysical properties such as diffusion constants and anomalous exponents are determined for each pixel to create two dimensional maps of chromatin dynamics at single pixel resolution in living single cells. Hi-D created spatially resolved maps show that DNA compaction and dynamics do not necessarily correlate. Instead, the maps suggest that chromatin dynamics are dictated by DNA-DNA contacts and protein binding to DNA, rather than chromatin density.

## RESULTS

### Hi-D maps genome dynamic properties at nanoscale resolution in living cells

Motion of densely distributed fluorescent molecules was quantitatively reconstructed from a series of conventional confocal fluorescence microscopy images by a dense Optical Flow method [20]. By integrating the resulting flow fields, a trajectory was computed for each pixel (Fig 1a; Additional file 1: Note S1; Additional file 1: Fig S1). The type of diffusion characterizing each pixel’s chromatin motion was chosen in an unbiased manner using a Bayesian inference from a set of five common models to fit each trajectory’s Mean Squared Displacement (MSD) [21] (Fig 1b, left panel; Additional file 1: Fig S2). The best fitting models were directly mapped onto the nucleus (Fig 1b; right panel) (Methods section). We found that only a small fraction of trajectories displayed directed diffusion (Fig 1b), while the bulk of chromatin exhibited sub-diffusive behaviour. Distinguishing between the comparable cases of anomalous and confined diffusion is a challenging task, given the limited duration of the experiment. After examination of a range of parameters governing these different types of diffusion, our results suggest that chromatin diffusion in human U2OS cells can be adequately described as anomalous to avoid misinterpretation (Additional file 1: Note S2; Additional file 1: Fig S3). The resulting biophysical parameters calculated for each pixel by Hi-D (diffusion constant *D*, anomalous exponent *α* and drift velocity *V*) are presented in color-coded 2D heat-maps (Fig 1c) (Methods section). They are distributed in a mosaic of irregular shape and dimensions of similar values (Fig 1c). These parameter maps clearly demonstrate that chromatin dynamics are spatially heterogeneous and partitioned. These maps also illustrate the notion that chromatin dynamics are spatially correlated in the micrometer-range [18,20]. To further characterize this heterogeneous distribution, the parameter distributions were deconvolved into discrete sub-populations using a General Mixture model (GMM) (Fig 1c; Methods; and Additional file 1: Fig S4). The GMM identified three populations of chromatin mobility referred to as slow, intermediate and fast (Methods; exemplary in Fig 1c), irrespective of the parameter under consideration (diffusion constant or anomalous exponent) or transcriptional state of the cell. We found that chromatin dynamics characterized by directed motion involving a drift velocity (V) was less present than free and anomalous diffusion and provided significantly less data for V than for the other two parameters (Fig 1c). Hence, drift velocity was not retained for further analysis.

**Fig 1:**
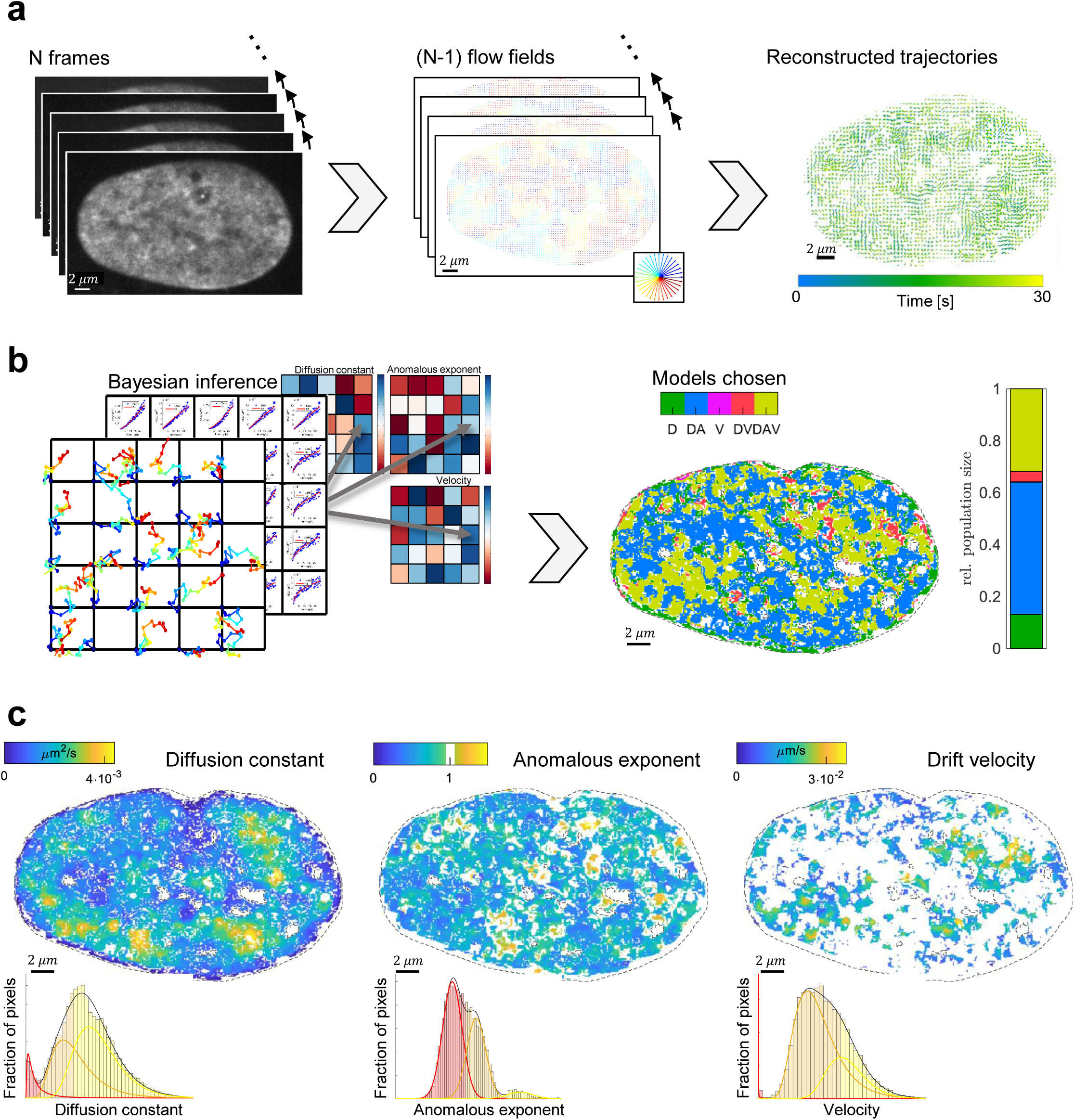
Hi-D enables spatially resolved mapping of genome dynamic properties at nanoscale resolution in living cells. Workflow: **a)** A series of N=150 confocal microscopy images acquired at 5 fps (left) (here SiR DNA stained living U2OS cells). Dense optical flow was applied to define (N-1) flow fields of the images (center, color coded) based on fluorescence intensity of each pixel (size = 65 nm). Individual trajectories are reconstructed over the duration of acquisition (right). **b)** MSD model selection (left): Trajectories of a 3×3 neighborhood of every pixel are used to calculate a mean MSD curve and its corresponding covariance matrix. By a Bayesian inference approach, the type of diffusion fitting each individual curve is chosen (free diffusion (D), anomalous diffusion (DA), directed motion (V) or a combination (DV) or (DAV). The spatial distribution of the selected models for each pixel is shown as a color map. **c)** Maps of biophysical parameters (*D, α* and *V*) extracted from the best describing model per pixel reveal local dynamic behavior of DNA in large domains. The distribution is deconvolved using a General Mixture Model.

### Validation of the Hi-D approach in simulation and experiment

In order to examine the suitability of calculated trajectories and associated diffusion constants by Hi-D to whole-chromatin imaging conditions, we compared Hi-D to dynamic multiple-target tracing (MTT), a Single Particle Tracking (SPT) method which is commonly used for dense molecule tracking [22] (Fig 2a,b; Additional file 1: Note S3; Additional file 1: Fig S5; Additional file 1: Fig S7). While the SPT method outperforms the Hi-D approach in scenarios of sparsely labelled molecules (Fig 2a), Hi-D analysis provided considerably more accurate estimates of local diffusion constants than SPT in scenarios of densely labelled molecules or structures of heterogeneous label density such as chromatin (Fig 2b). Hi-D therefore constitutes an approach to extract dynamic parameters of biomolecules with dense labeling where SPT is unsuitable. One should, however, keep in mind that SPT and Hi-D are meant to analyze images from drastically different labeling conditions and should thus refrain from a direct comparison between single-locus dynamics analyzed by SPT and local bulk chromatin dynamics by Hi-D (Additional file 1: Note S4; Additional file 1: Movie S1).

**Fig 2:**
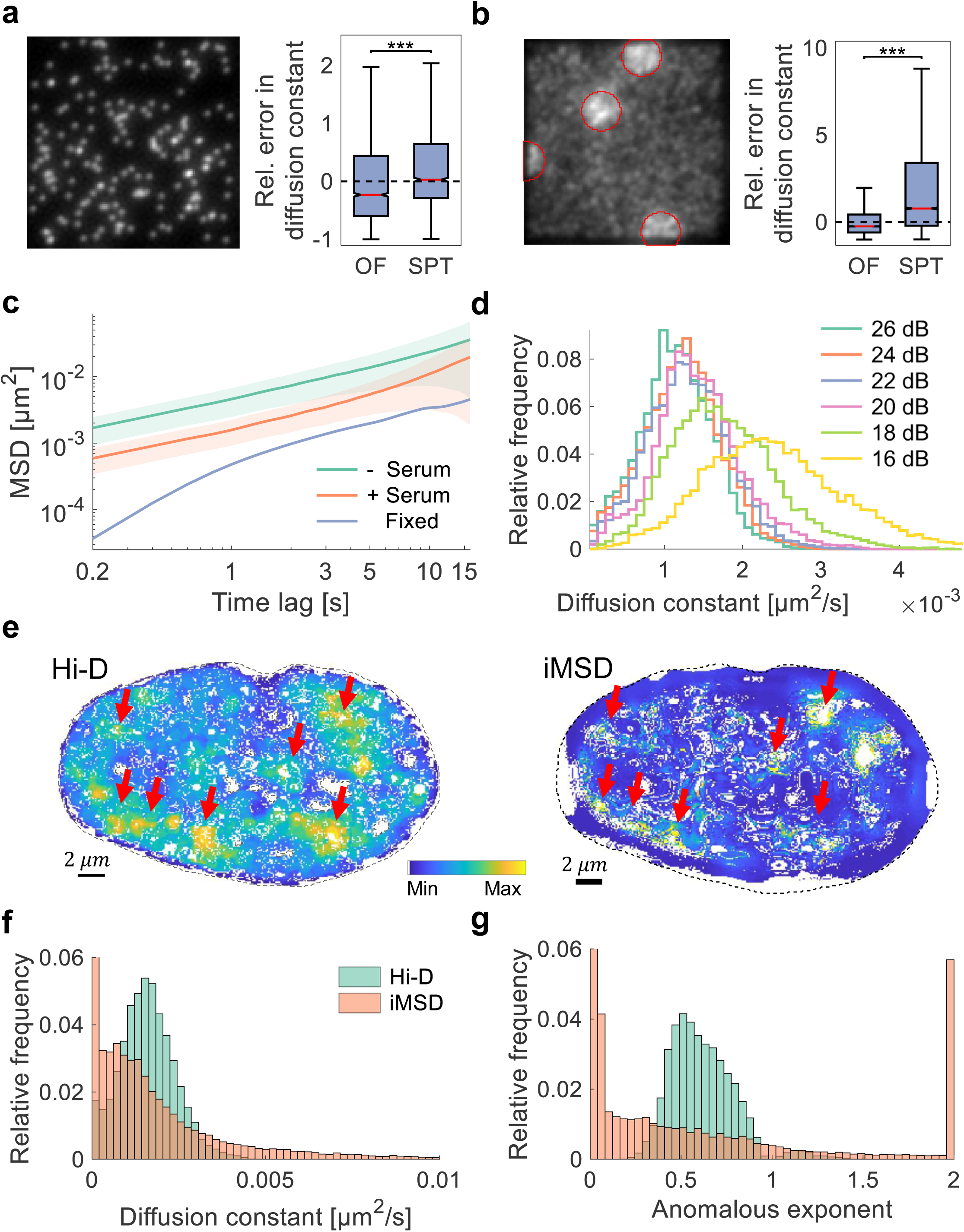
Experimental validation of the Hi-D approach. **a)** Exemplary frame of a simulated time series with low density (0.001/*px*^3^) of emitters undergoing Brownian motion convolved by a typical point spread function (left). The time series is subject to Hi-D and Single Particle Tracking estimating the trajectories of emitters. From the estimated trajectories, the MSD is computed and compared to the ground truth diffusion constant. The relative error in the determined diffusion constant is shown. **b)** High density (0.02/*px*^3^) of emitters with patches of super-high density (0.035/*px*^3^) encircled for visualization, imitating regions of densely packed chromatin. Dashed lines show the optimal value, i.e. perfect agreement between estimation and ground truth. Red lines indicate the median value. Data from 10 independent simulations. Statistical significance assessed by a two-sample Kolmogorov-Smirnov test (***: p < 0.001). **c)** MSD curves computed in fixed (n = 13) and living quiescent (- serum; n = 13) and serum stimulated (+ serum; n = 14) U2OS cells. Diffusion constants for the three average curves were derived by regression yielding *D* = (0.87 ± 0.1) · 10^−3^ *μm*^2^/*s* for quiescent, *D* = (2.6 ± 0.1) · 10^−4^ *μm*^2^/*s* for stimulated and *D* = (6.1 ± 0.1) · 10^−6^ *μm*^2^/*s* for fixed cells. MSD curves show considerably higher MSD values for living cells and diffusion constants are two orders of magnitude higher for living cells thus confirming the detection of motion well above noise background. **d)** Diffusion constants derived from a nucleus corrupted with varying levels of signal-to-noise ratio. Results are consistent up to a lower bound of ∼ 20 dB. **e)** Map of diffusion constants computed by Hi-D (left) and iMSD (right). Diffusion constants are color-coded from their minimum to their maximum value (blue to yellow; for absolute values see f)). Red arrows indicate regions of high mobility detected by both methods. **f)** Diffusion constants shown in e) and **g)** corresponding values of the anomalous exponent computed by Hi-D (blue) and iMSD (red).

To ensure that the calculated dynamics are not a consequence of imaging noise, we experimentally validated the sensitivity of the approach by calculating the MSD for formaldehyde fixed and living U2OS cells labelled by DNA-SiR in quiescence (- serum) or normal growth (+ serum). Diffusion constants derived from the MSD curves by Bayesian inference were about two orders of magnitude greater in living cells than in fixed cells (Fig 2c) confirming that Hi-D enables quantifying DNA dynamics well above the noise background. To confirm the robustness of extracted parameter values with respect to varying levels of imaging noise, Hi-D was applied to nuclei to which noise was artificially added. The signal-to-noise ratio (SNR) of the original nuclei were about 26 dB and subsequently reduced stepwise down to 16 dB. The distributions of computed diffusion constants were consistent up to a lower limit of about 20 dB, below which the distribution is considerably biased towards larger values and broadens (Fig 2d, Additional file 1: Fig S8). Likewise, features of the spatial map of diffusion constants were equally conserved for SNR values as low as 20 dB, demonstrating the robustness of Hi-D for varying imaging noise (Additional file 1: Fig S8). In analogy to the robustness to varying SNR levels, Hi-D is thus robust to photobleaching effects (if SNR ≥ 20dB) since flow fields are only estimated between consecutive images, for which illumination changes due to photobleaching are usually negligible. Furthermore, Hi-D was also shown to be robust to small variations in time intervals of acquired time series as long as the expected motion between frames was in the order of the pixel size (Additional file 1: S9). We further validated Hi-D against iMSD, a well-established method to extract dynamic information of dense molecules, based on the spatial correlation function of intensity fluctuations caused by diffusing molecules, which are recorded using camera-based systems [23]. Using successive calculations of iMSD to overlap regions of interest, we computed a diffusion map similar to Hi-D derived maps (Additional file 1: Note S5). Quantitatively, both methods yield diffusion constants of the same order of magnitude (Hi-D (1.6 ± 0.8) · 10^−3^ *μm*^2^/*s*, iMSD (2.2 ± 4.5) · 10^−3^ *μm*^2^/*s*, mean ± standard deviation), which are consistent with reported values using SPT and correlation spectroscopy methods applied to interphase chromatin [14,18]. However, the distribution of values derived by iMSD was considerably broader than the distribution revealed by Hi-D (Fig 2f). The distribution of anomalous exponents computed by iMSD showed many spurious values at the limit of the scale, while Hi-D consistently returns reasonable values (Fig 2g). We thus conclude that Hi-D reveals dynamic parameters of the same order of magnitude as iMSD but is advantageous in the estimation of multiple parameters simultaneously, by virtue of the featured Bayesian model selection. Hi-D is thus an accurate, robust, fast and easy to use tool to determine dynamics of macromolecules nucleus-wide.

### Single-cell biophysical property maps of genome conformation and behaviour

To concomitantly monitor position and distribution of the DNA mobility populations under different biological conditions, we determined Hi-D maps of the same serum-starved and then stimulated cell (Fig 3a). Transcription is largely inhibited in cell-cycle arrested cells grown in a serum-free medium (Additional file 1: Fig S10). Adding serum to the medium stimulates mRNA production through transcriptional activity [20,24–26]. As above, diffusion constants of DNA motion were calculated for each pixel based on the model selected by Bayesian inference. Small diffusion constants characterized motion of chromatin prominently located at the nuclear envelope (dark blue). Plotting the average diffusion constants versus the distance from the nuclear periphery showed that the mobility within a rim of 1 µm from the periphery increases linearly before adopting a nearly constant value in the inner volume of the nucleus (Additional file 1: Fig S11). At numerous sites across the remaining nuclear volume, fast diffusive areas of irregular dimensions spanning 0.3 − 3 *μm* in diameter (yellow areas in Fig 3a) are embedded in the bulk of moderately dynamic chromatin. Areas of different parameter values seamlessly transition into one another without clearly defined boundaries, reminiscent of spatially correlated chromatin dynamics [20]. Upon serum stimulation, the spatial distribution of high and low diffusion constants was largely conserved (compare the presence of yellow regions in the quiescent and serum stimulated cell in Fig 3a), but the diffusion constant was globally strongly reduced by nearly one order of magnitude. Deconvolution of the distribution of diffusion constants and labelling of pixels according to the mobility population determined by the GMM (Fig 3b; slow - red, intermediate – orange and fast - yellow) yields a map in agreement with this observation. Deconvolution hence classifies regions according to the values of a given parameter compared to other regions within the same nucleus. When nuclear activity is modulated, changes in this classification can be measured. In particular, the fast diffusing population with respect to the bulk chromatin in quiescent cells is reduced upon serum stimulation (Fig 3c) and re-classified as intermediate population. Connected areas with high mobility appear eroded (Fig 3b). In contrast, the slow population occupying ∼ 6 % of the nuclear area, which is almost exclusively located at the nuclear periphery, was invariant to transcriptional changes (Fig 3b, c). Despite considerable reorganization of the relative distribution of mobility populations and overall reduced intensity of motion, the type of diffusion governing the nuclear parameter maps showed only moderate changes upon stimulation of transcriptional activity (Fig 3d).

**Fig 3:**
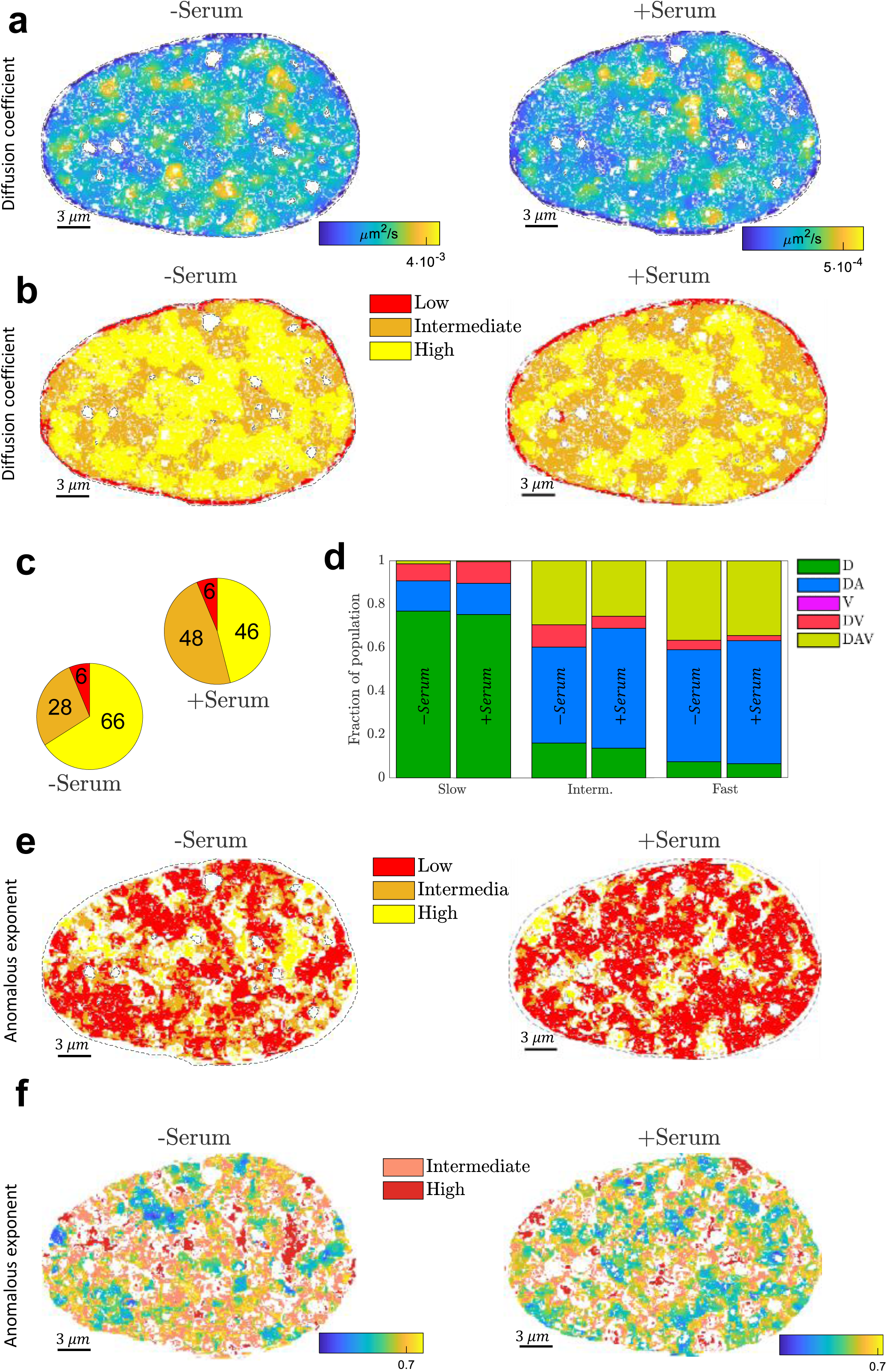
Hi-D maps single-cell biophysical properties of genome conformation and behaviour. **a)** Diffusion constant spatially mapped onto the nucleus of a quiescent cell (left) and the same nucleus upon serum stimulation **b)** Spatial distribution of populations found by the GMM for the diffusion constant for quiescent (left) and actively transcribing cells (right). **c)** Relative share of populations on the cell volume (n = 12). Numbers in percent. **d)** For each population in b), the relative share of chosen MSD models is represented as a stacked histogram in the quiescent state (left bars) and actively transcribing state (right bars). **e)** Spatial distribution of populations for the anomalous exponent for quiescent (left) and actively transcribing cells (right). **f)** Detailed insight into the spatial patterning of the low population of the anomalous exponent. The intermediate and high population are shown in light and dark red respectively.

Anomalous diffusion dominated across the entire nucleus (0.3 ≤ *α* ≤ 0.73) forming a mosaic-like pattern, which underwent, compared to maps of diffusion constants, considerable remodeling upon transcriptional activation (Fig 3e, f). Within this pattern, patches of super-diffusive (red: *α* > 1) motion segregated into distinct islands which became more fragmented upon serum stimulation. Random contacts or re-distribution of existing contacts of the chromatin with itself may give rise to such variations in anomalous exponent upon serum stimulation [27]. Because the diffusion constant of chromatin fibers appears unaffected for moderate degrees of cross-linking [28], we expect that association of proteins with DNA upon serum stimulation could favor global decrease of mobility in vivo. Hi-D reveals high-resolution spatial changes in mobility and in anomaly of chromatin diffusion in single cells. Further investigation may tell us if all or a subset of visible physical domains correspond to the ones determined using chromosome conformation capture (Hi-C).

### Transcription modulates chromatin and RNA polymerase II motion

To further explore the relationship between global chromatin dynamics and transcriptional activity, we examined the dynamics of RNA polymerase II (RPB1-Dendra2; RNA Pol II) in live U2OS cell nuclei (Fig 4a) at different transcriptional states. Hi-D analysis resolved three mobility populations of RNA Pol II (Fig 4b), which is consistent with the existence of three kinetically different groups of RNA Pol II based on the half-life of chromatin-binding [29,30]. Diffusion constants of the three dynamic populations in actively transcribing cells (grown in normal condition) were significantly greater compared to transcriptionally less active cells (serum-starved cells) (Fig 4b). In quiescent cells, the fraction of quickly diffusing RNA Pol II complexes was reduced compared to actively transcribing cells. Upon elongation inhibition using 5,6-Dichloro-1-β-D-ribofuranosylbenzimidazole (DRB), the slowly diffusing fraction was greater than in untreated cells, indicating tenacious immobilization of RNA Pol II on the DNA template after initiation (Fig 4c). The average diffusion constants in serum starved and DRB treated cells stayed roughly unchanged in all three populations, suggesting that RNA Pol II is unbound in the absence of serum [25].

**Fig 4:**
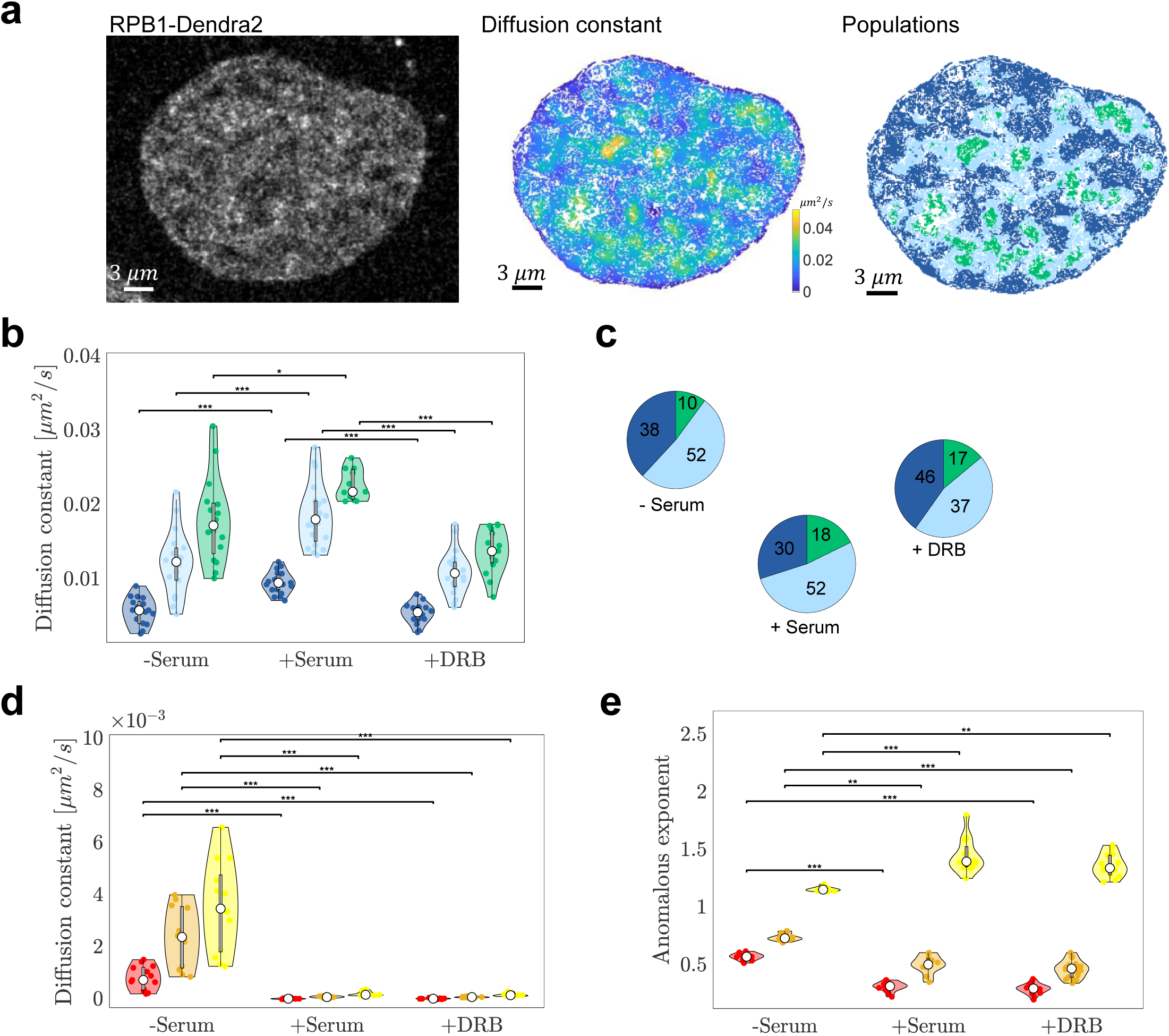
Biophysical properties of chromatin are sensitive to transcriptional activity. **a)** Fluorescence image of RPB1, a RNA polymerase II subunit, fused to Dendra2 (left), the spatial mapping of the diffusion constant (middle) and population deconvolution (right). **b)** Violin plots of the mean diffusion constant of RNA Pol II for all three mobility groups in quiescent (- serum; n = 18), actively transcribing (+ serum; n = 20) and elongation inhibited (+ DRB; n = 21) cells are shown; dark blue, light blue and green denote the slow, intermediate and fast population, respectively. **c)** Relative share of the populations on the cell volume for starved (n = 13), stimulated (n = 14) and DRB-treated (n = 18) cells. Numbers in percent. **d)** As b) for the diffusion constant and **d)** the anomalous exponent of DNA dynamics (n=13 cells); red, gold and yellow denote the slow, intermediate and fast population, respectively. Statistical significance for assessed by a Friedman test (*: p < 0.05, **: p < 0.01, ***: p < 0.001).

We then compared the effect of transcriptional activity on chromatin dynamics in serum-starved and –stimulated cells. In contrast to RNA Pol II mobility, the average diffusion constant of DNA in serum-starved U2OS nuclei decreased by nearly one order of magnitude for all three populations upon addition of serum. Arresting RNA Pol II before elongation did not change the observed diffusion constants, compared to undisturbed transcription (Fig 4d). These cell-population results are consistent with results from single-cell analyses (Fig 3) and strengthen our hypothesis that nuclear processes considerably hamper diffusion of chromatin. In a quiescent state, only essential nuclear processes are maintained. Fewer protein complexes acting upon DNA in a could facilitate motion of the chromatin fiber. Upon serum stimulation, binding of transcription factor complexes and other proteins to DNA increase crowding, reduce the freedom to move and hence the apparent chromatin dynamics, at least in a subset of domains. Increased DNA-protein interactions and interchromatin contacts also enhance spatial correlation of chromatin dynamics in serum supplemented compared to quiescent cells [20]. Serum addition to starved cells likely stimulated RNA Pol II binding to DNA. When inhibiting elongation, transcription factories are still present [24] and, in agreement with chromatin coherence, DNA mobility remains constrained [20].

Independently of the culture conditions, a ground-state Rouse-like behavior characterizes chromatin in the examined nuclei (MSD fit with *α* close to 0.5) [31,32]. Upon serum stimulation of starved cells, anomalous diffusion became predominant and its value (*α* ∼ 0.33) is indicative of entangled polymers [33]. This behavior was also determined for a single labelled site next to an actively transcribing gene [13]. Entanglement could stem from random DNA-protein contacts, a model coherent with polymer simulations inspired by chromosomal capture data [34]. Hindered motion of chromatin and RNA Pol II is thus a direct consequence of forming transcription ‘hubs’ or factories to which chromatin is tethered [25].

### Chromatin dynamics is uncoupled from compaction

We next asked if chromatin dynamics are influenced by the compaction of chromatin since heterochromatin is widely believed to be less dynamic than euchromatin [16]. Eu- and heterochromatin domains were determined in serum-starved and –stimulated cells by quantifying fluorescence intensity as described in [35] (Fig 5a). We found that the average flow magnitude between successive frames was independent of the compaction state of chromatin (Fig 5b). Likewise, the distribution of diffusion constants did not correlate with chromatin density or euchromatin and heterochromatin (Fig 5c). Peripheral heterochromatin overlapped with the slow motion domain at the nuclear rim (Fig 5d) consistent with previous findings [16]. In the inner nuclear volume however, we did not observe any tendency of heterochromatin being associated with a specific mobility population. Instead, we observed that mobility populations were distributed randomly among euchromatin and heterochromatin regions (Fig 5e) with the exception that in serum-starved cells, the heterochromatin fraction was slightly enriched in the slow diffusing population at the nuclear periphery. Furthermore, we found that regions characterized by a specific anomalous exponent did not preferentially overlap with either eu- or heterochromatin (Fig 5f, g). These results also hold for MCF7 cells and different fluorescent markers for chromatin (Additional file 1: Fig S12). These findings were confirmed in NIH3T3 cells expressing GFP-HP1α, a well-established marker for heterochromatin (Additional file 1: Fig S13). In addition, Hi-D analysis of HP1α hints towards previously proposed liquid phase-separation [36]. Our results thus suggest that chromatin undergoes diffusion processes which are, in general, unrelated to the compaction level of chromatin. However, compact chromatin is characterized by increased contact frequency of the chromatin fiber with itself, which could enhance the extent of coherent chromatin motion. To test this hypothesis, we calculated Moran’s Index of Spatial Autocorrelation [37] for the flow magnitude assessed at different time lags in eu- or heterochromatin (Fig 5h). We found that heterochromatin exhibits enhanced spatial autocorrelation compared to euchromatin across all accessible time lags. Furthermore, the spatial autocorrelation decreases with increasing time lags in serum-starved cells, while in serum-stimulated cells, autocorrelation is enhanced in the long-time limit (over 30 seconds). This finding points to active processes establishing spatial coherence in the long-term [18,20] while random processes such as thermal fluctuations decrease autocorrelation at time scales greater than 10 seconds in serum-starved cells.

**Fig 5:**
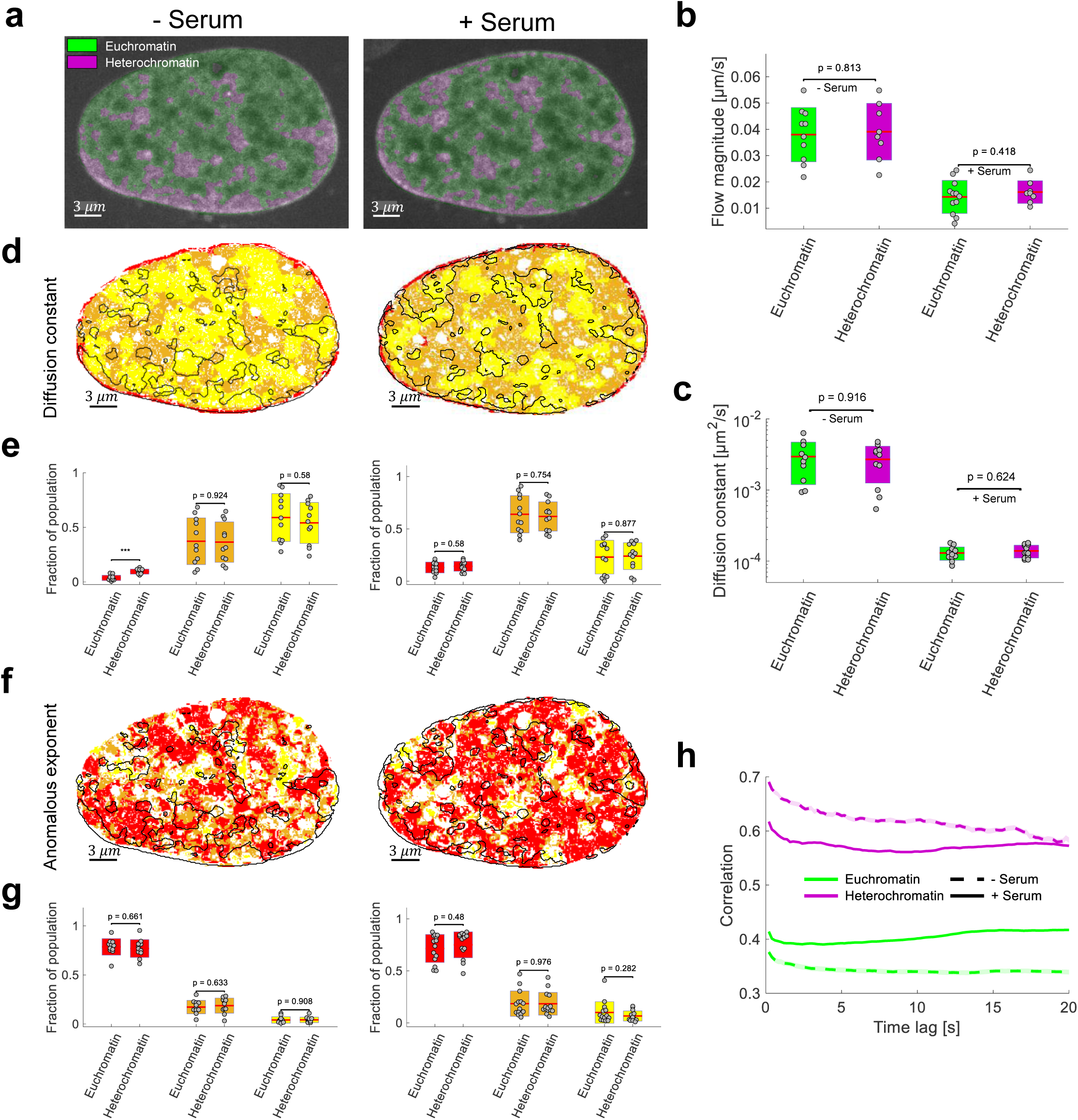
Chromatin compaction and dynamics do not spatially correlate. **a)** Spatial classification of signal intensity into euchromatin and heterochromatin [35] overlaid on an exemplary fluorescence image for quiescent (left) and stimulated (right) cells. **b)** Average flow magnitude and **c)** diffusion constant (n = 12) in euchromatin and heterochromatin for starved (left) and serum stimulated cells (right). Statistical significance assessed by a two-sample t-test. **d)** Overlay with the diffusion populations found by Hi-D. Black solid line corresponds to eu- / heterochromatin region boundaries. **e)** Diffusion populations show a similar distribution over hetero- and euchromatin. The colors refer to the slow, intermediate and high population respectively and each point corresponds to one nucleus. Statistical significance assessed by a two-sample t-test (*: p < 0.05, **: p < 0.01, ***: p < 0.001). **f-g)** Anomalous exponent as d-e). **h)** Spatial autocorrelation at euchromatin (green) and heterochromatin (purple) of the flow magnitude between all accessible time lags in quiescent and serum-stimulated cells.

## DISCUSSION

Hi-D analysing the dynamics of dense structures such as chromatin and RNA Pol II directly in single cells without losing active fluorophore density and with no need for prior experience in sophisticated labelling preparations or advanced microscopy [38]. We show that Hi-D is an accurate and robust read-out of chromatin dynamics and that the information gain through image analysis afforded by Hi-D alleviates the incompatibility of conventional microscopy for nanoscale mapping of properties of nuclear dynamics in living cells.

Hi-D analysis revealed that DNA dynamics can roughly be classified into three sub-populations within the mammalian nucleus. The use of three distinct populations does not reflect real, likely fluctuating transitions between the populations, but is a handy means to characterize the highly complex and heterogeneous dynamic landscape of chromatin. The first population, a slow mobility fraction prominently located at the nuclear rim, is reminiscent of lamina associated domains (LADs) [39]. The dynamic response of this population to transcriptional stimuli supports the hypothesis that LADs play an important role in attenuating transcription activity and in controlling gene expression [40]. Although less mobile chromatin at the nuclear periphery largely overlapped with long known perinuclear heterochromatin [41,42], Hi-D analysis remarkably points to an overall absence of correlation between chromatin compaction and mobility. Indeed, two non-peripheral chromatin subpopulations which display intermediate and highly diffusive regimes, are distributed in a mosaic-like pattern throughout the nucleus and include sections of intranuclear heterochromatin. Heterochromatin therefore does not exhibit low mobility in nuclear space in general, but may be divided into a more viscous component with reduced mobility due to anchoring to a nuclear lamina and a more rigid LAD component tethered to nuclear structures [39,43–45]. These results are coherent with the hypothesis that heterochromatin domains are formed by a liquid phase-separation mechanism characteristic of mobility reduction across phase boundaries [36]. The extent of the third, highly mobile fraction, which dominated in the quiescent state, decreased dramatically when cells were serum stimulated. This switch in chromatin mobility is suggestive of altered DNA-protein interactions and accrued local concentration of proteins in a highly transcribing nucleus [25].

Heterogeneous chromatin motion arises due to irregular protein binding along the chromosomes and can lead to thermodynamic or electrostatic self-organisation of nuclear compartments [46,47]. Local patches of large anomalous exponents indicate super-diffusive behaviour of chromatin which may result, among others, from active noise acting on the chromatin fiber [48,49], even in quiescent cells. Chromatin patches with *α* < 0.3 and *α* > 1 respectively correspond in size to one or a few DNA loops [50]. These two types of patches are present as islands within the general chromatin fraction governed by an anomalous exponent 0.3 < *α* < 1. This organization is also in good agreement with the chromosome territory – interchromatin compartment model [2]. In conclusion, the combination of diffusion constant and anomalous exponent maps provides an integrated view on chromatin dynamics and yields insights into possible mechanisms driving dynamics as well as the local chromosomal organization and re-organization during genomic processes.

Our results support the hypothesis that a short treatment of DRB is sufficient to halt most of a cell’s transcription [51], but it is believed to have a limited effect on chromatin configuration of promoters, or on upstream events and assembly dynamics of the preinitiation complex (PIC) [52]. Our results suggest that a few minutes of DRB treatment affects only the assembly dynamics of the RNA Pol II itself. In contrast, longer treatment with DRB as well as prolonged serum starvation, are expected to result in massive PIC disassembly. We believe therefore that increased chromatin mobility upon serum starvation correlates with PIC breakdown. Conversely, when serum is present in the medium, activity of RNA Pol II and of PIC enzymes is restored and PIC will assemble onto DNA, both pervasively across the genome at background levels (pervasive transcription) and more stably at genes [53,54]. Reduced mobility of chromatin in the presence of serum suggests that stable PIC binding may serve as an anchoring function for individual chromatin fibres, on top of its essential function in transcription initiation.

Hi-D can be applied to real-time imaging of any abundant fluorescent molecule to obtain comprehensive maps of their dynamic behaviour in response to stimuli, inhibitors or disruptors of nuclear functions and integrity.

## METHODS

### Cell Culture

A Human U2OS osterosarcoma cell line (for DNA imaging) and MCF-7 cells (ATCC) were maintained in Dulbecco’s modified Eagle’s medium (DMEM) containing phenol red-free and DMEM-12 (Sigma-Aldrich), respectively. For RNA Pol II imaging, a U2OS cell line stably expressing RPB1 (subunit of RNA Pol II) fused with Dendra2 was constructed as previously described in [55]. Medium was supplemented with Glutamax containing 50 μg/ml gentamicin (Sigma-Aldrich), 10% Fetal bovine serum (FBS), 1 mM sodium pyruvate (Sigma-Aldrich) and G418 0.5 mg/ml (Sigma-Aldrich) at 37°C with 5% CO_2_. Cells were plated for 24 h on 35 mm petri dishes with a #1.5 coverslip like bottom (μ-Dish, Ibidi, Biovalley) with a density of about 10^5^ cells/dish.

### DNA staining

U2OS and MCF-7 cell lines were labelled by using SiR-DNA (SiR-Hoechst) kit (Spirochrome AG). For DNA was labelled as described in [56]. Briefly, we diluted 1 mM stock solution in cell culture medium to concentration of 2 μM and vortexed briefly. On the day of the imaging, the culture medium was changed to medium containing SiR-fluorophores and incubated at 37°C for 30-60 minutes. Before imaging, the medium was changed to L-15 medium (Liebovitz’s, Gibco) for live imaging.

### Cell starvation and stimulation

For starvation mode, cells were incubated for 24 h at 37°C before imaging with serum-free medium (DMEM, Glutamax containing 50 μg/ml gentamicin, 1 mM sodium pyruvate, and G418 0.5 mg/ml). Just before imaging, cells were mounted in L-15 medium. For stimulation, 10% FBS was added to the L-15 medium for 10-15 minutes.

### Flow cytometry analysis with Hoechst and Pyronin Y staining

To differentiate cells in G0 versus G1, double staining of Hoechst 33324 and Pyronin Y was used, as previously described [26,57]. Briefly, U2OS cells representing each condition (with serum, without serum during 24h and without serum during 24h + 15 min serum) were trypsinized, cells were collected into phosphate-buffer saline (PBS) at a concentration of 2×10^6^ cells/ml, then added to a fixative of ice-cold 70% ethanol. Cells were fixed for at least 2h and washed twice with FACS buffer (1xPBS supplemented with 2% (v/v) heat-inactivated, sterile-filtered fatal bovine serum, 1 mM EDTA). Cells were then incubated in a water bath pre-adjusted to 37°C for 45 min in the dark with 2 µg/ml of Hoechst 33324 (Invitrogen, H3570) diluted in FACS buffer, then incubated to 37°C for 30 min in the dark with 4 µg/ml of Pyronin Y (abcam, ab146350), diluted in FACS buffer. The samples were kept in the dark at 4°C until analyzed using a LSR II flow cytometer (BD Biosciences). Hoechst 33342 and Pyronin Y staining were measured with a UV (350 nm) and yellow green (561 nm) lasers, respectively. DNA content was determined by Hoechst 33342 and RNA content was determined by Pyronin Y. Cells in G0 were identified as the population with 2N DNA content and RNA content lower than the level of cells assigned to G1 phase. Medians of Pyronin Y staining for each condition were compared to assess the increase in mRNA production after serum addition.

### DRB treatment

The U2OS cells for both DNA and RNA Pol II images were treated with 100 μM 5,6 Dichlorobenzimidazole 1-β-D-ribofuranoside (DRB; Sigma-Aldrich) for transcription inhibition prior to live-cell image acquisition. DRB was diluted in the L-15 (Leibovitz) imaging medium that was supplemented with 10% FBS, DMEM, Glutamax containing 50 μg/ml gentamicin, 1 mM sodium pyruvate, and G418 0.5 mg/ml. The imaging medium was changed with fresh L-15 medium containing DRB and incubated under the microscope for 15 min before imaging.

### Cell fixation

U2OS cells were washed with a pre-warmed (37 °C) phosphate buffered saline (PBS) and followed by fixation with 4% (vol/vol) Paraformaldehyde in PBS for 10-20 min at room temperature. Images were recorded at room temperature in PBS, after washing the cells with PBS (three times, 5 min each).

### DNA live cell imaging

Cells were placed in a 37 °C humid incubator by controlling the temperature and CO_2_ flow using H201-couple with temperature and CO_2_ units. Live chromatin imaging was acquired using a DMI8 inverted automated microscope (Leica Microsystems) featuring a confocal spinning disk unit (CSU-X1-M1N, Yokogawa). An integrated laser engine (ILE 400, Andor) was used for excitation with a selected wavelength of 647 nm and 140 mW as excitation power. A 100x oil immersion objective (Leica HCX-PL-APO) with a 1.4 NA was chosen for a high-resolution imaging. Fluorescence emission of the SiR–Hoechst was filtered by a single-band bandpass filter (FF01-650/13-25, Semrock, Inc.). Image series of 150 frames (5 fps), with exposure time of 150ms per frame were acquired using Metamorph software (Molecular Devices) and detected using sCMOS cameras (ORCA-Flash4.0 V2) and (1×1 binning), with sample pixel size of 65 nm. All series were recorded at 37°C.

### RNA Pol II live cell imaging

Image series of 150 frames were recorded with an exposure time of 200 ms using a Nipkow-disk confocal system (Revolution, Andor) featuring a confocal spinning disk unit (CSU22, Yokogawa). A diode-pumped solid-state laser with a single wavelength of a 488 nm (Coherent) at 5-10% laser power was used for excitation of RPB1 fused to Dendra2. A 100x an oil immersion objective (Plan Apo 1.42, Nikon) was used for imaging. The fluorescent emission signal was filtered through an emission filter (ET525/30-25, Semrock, Inc.), and detected at 512/18 nm on a cooled electron multiplying charge-coupled device camera (iXon Ultra 888), with sample pixel size of 88 nm.

### Image processing

#### Denoising

Raw images were denoised using non-iterative bilateral filtering [58]. While Gaussian blurring only accounts for the spatial distance of a pixel and its neighbourhood, bilateral filtering additionally takes the difference in intensity values into account and is therefore an edge-preserving method. Abrupt transitions from high- to low-intensity regions (e.g. heterochromatin to euchromatin) are not over-smoothed. Images of varying noise levels were treated with a bilateral filter with half-size of the Gaussian bilateral filter window of 5 pixels, the spatial-domain standard deviation value was set to 5 pixels and the intensity-domain standard deviation was varied from 0.3 to 0.8 for decreasing levels of the signal-to-noise ratio from 26 dB to 16 dB (*SNR* = 10 log_10_(*I*^2^/*σ*^2^), with the signal power *I*^2^ and the noise variance *σ*^2^).

### MSD analysis and model selection by using Bayesian inference

In order to carry out a MSD analysis locally, the spatial dependency of the Mean Squared Displacement (MSD) can be written explicitly:

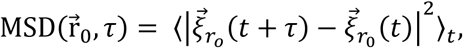

where 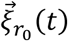 is the position at time *t* of a virtual particle with initial position 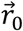, *τ* = {Δ*t*, 2Δ*t*, …, (*N* − 1)Δ*t*} are time lags where Δ*t* is the time difference between subsequent images and the average <·>_*t*_ is taken over time. The resulting MSD is a function of the initial position 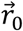 and the time lag *τ*.

#### MSD models

The MSD can be expressed analytically for anomalous diffusion (DA), confined diffusion (DR) and directed motion (V) in two dimensions as

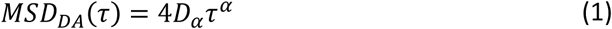

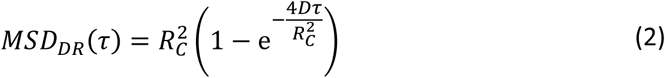

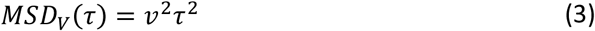

where *D*_*α*_ is the diffusion constant in units of *μm*^2^/*s*^*α*^, *α* is its anomalous exponent, *v* [*μm*/*s*] its velocity and *R*_*C*_ [*μm*] is the radius of a sphere within the particle is confined [59]. The case *α* = 1 is known as free diffusion, 0 < *α* < 1 corresponds to anomalous diffusion and 1 < *α* ≤ 2 corresponds to superdiffusion. Strictly speaking, each generalized diffusion constant *D*_*α*_ has different units, corresponding to the specific value of *α*. However, we refer to it as the diffusion constant *D* throughout the text for simplicity. Additionally to eq. (1)-(3), different types of motion can appear overlaying, resulting in a linear combination of the equations above. For example, anomalous motion can be superimposed on an underlying drift and the resulting *MSD* reads *MSD*_*DAV*_ (*τ*) = *MSD*_*DA*_(*τ*) + *MSD*_*V*_ (*τ*). We found that anomalous and confined diffusion appears very similar in experimental data and therefore decided in favor for anomalous diffusion to describe our data (Additional file 1: Note S3). The abbreviations used in this study are summarized in Table 1. As experimental data is usually subject to noise, a constant offset *o* is added to every model.

**Table 1:** Overview over possible Mean Squared Displacement models.

| Abbreviation | Model | Formula |
| --- | --- | --- |
| D | Free diffusion | $MSD_D(\tau) = 4D\tau + o$ |
| DA | Anomalous diffusion | $MSD_{DA}(\tau) = 4D_\alpha \tau^\alpha + o$ |
| V | Drift | $MSD_V(\tau) = v^2 \tau^2 + o$ |
| DV | Free diffusion + drift | $MSD_{DV}(\tau) = 4D\tau + v^2 \tau^2 + o$ |
| DAV | Anomalous diffusion + drift | $MSD_{DAV}(\tau) = 4D_\alpha \tau^\alpha + v^2 \tau^2 + o$ |

#### MSD model selection

The MSD is calculated for every pixel independently, resulting in a space- and time lag-dependent MSD. It is known that living cells can behave largely heterogeneous [3,60]. Ad-hoc, it is not known which diffusion model is appropriate. Fitting an MSD curve with a wrong model might result in poor fits and highly inaccurate determination of the mentioned parameters. For this reason, we use a Bayesian inference approach to test different models for any given MSD curve as proposed by Monnier *et al*. [21]. Given the data *Y* = {*Y*_1_, …, *Y*_*n*_} and *K* model candidates *M* = {*M*_1_, …, *M*_*K*_}, each with its own (multidimensional) parameter set *θ* = {*θ*_1_, …, *θ*_*K*_}, we want to find the model *M*_*k*_(*Y, θ*_*k*_) such that the probability that *M*_*k*_(*Y, θ*_*k*_) describes the data, given the set of models to test, is maximal. By Bayes’ theorem, the probability for each model is given by

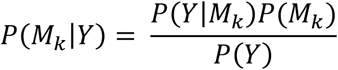

If there is no reason to prefer one model over the other, the prior probability of each model *P*(*M*_*k*_) is equal. The parameter set which is used to describe the data, given a fixed model, strongly influences the probability. Therefore, it is crucial to estimate the optimal parameters for every model in order to calculate the model probabilities. The probability that the data *Y* is observed, given the model *M*_*k*_ described by the model function *M*_*k*_(*x*; *θ*_*k*_) and any parameter set *θ*_*k*_ is approximated by a general multivariate Gaussian function [61]

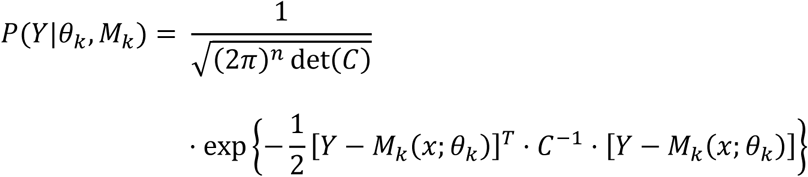

where *C* is the empirical covariance matrix of the data and the prefactor is a normalizing factor. This equation has an intuitive meaning. Assume we test a model *M*_*k*_ parametrized by *θ*_*k*_ to find out if it describes the data *Y*. The exponential function consists of the term [*Y* − *M*_*k*_(*x*; *θ*_*k*_)], i.e. the residuals of the data and the describing model. If the residuals are small, i.e. the model describes the data well, the exponent is small and the probability *P*(*Y*|*θ*_*k*_, *M*_*k*_) seeks 1. On the other hand, the worse the fit, the greater the resulting residuals and the probability seeks asymptotically to 0. The factor *C*^−1^ accounts for the covariance in the data. The covariance matrix for a set of MSD curves normally shows large values for large time lags as the uncertainty increases and MSD curves diverge. The covariance matrix implicitly introduces a weight to the data, which is small for large variances and large where the data spreads little. This fact avoids cutting of the MSD curve after a specific number of time lags, but instead includes all available time lags weighted by the covariance matrix. The approach is illustrated in (Additional file 1: Fig S2b) with the covariance matrix exemplary shown in the inset. In case of uncorrelated errors, non-diagonal elements are zero, but the approach keeps its validity [62] and follows an ordinary least-squares regression.

Given the best estimate of the parameter set for a model, the model and its corresponding parameters are chosen so that their probability to describe the data is maximal: 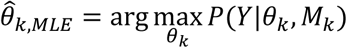. It has to be stressed that values of the anomalous exponent scatter around 1, but do not assume the value 1 (e.g. Fig 1c, middle panel). This is due to the model selection procedure, selecting the simplest model which is consistent with the data. In the case that the underlying motion is well described by free diffusion, *α* is inherently set to 1 and classified as free diffusion rather than anomalous diffusion. The descriptions of free diffusion or anomalous diffusion with *α* = 1 are equivalent, but the free diffusion model contains one parameter less and is therefore preferred leading to ‘missing’ *α* values close to 1 in the parameter maps and histograms. To carry out the MSD analysis locally, we choose to take the 3×3 neighborhood of a pixel, detect possible outliers therein by the interquartile range criterion [63] and calculate the error covariance matrix of the data within the pixel’s neighborhood. The restriction to a single pixel and its neighborhood allows us to carry out the MSD analysis of trajectories locally, in contrast to an ensemble MSD in previous studies [18], revealing only average information over many trajectories. The choice of a 3×3 window is reasonable with regard to the equivalently chosen filter size in the Optical Flow estimation. The flow field in this region is therefore assumed to be sufficiently smooth. All calculations, except for the General Mixture Model analysis, were carried out using MATLAB (MATLAB Release 2017a, The MathWorks, Inc., Natick, Massachusetts, United States) on a 64-bit Intel Xeon CPUE5-2609 1.90 GHz workstation with 64 GB RAM and running Microsoft Windows 10 Professional.

### Deconvolution of sub-populations

Regarding the distribution of diffusion constants, an analytical expression can be found assuming that the diffusion constant was calculated from a freely diffusing particle (*α* = 1)[64]. However, we find anomalous diffusion to a large extent in our data (e.g. Fig 1c, Fig 2g, Fig 3f, Fig 4e and Fig 5g) and, to our knowledge, an analytical expression cannot be found for distributions of anomalous exponent, radius of confinement and drift velocity. We therefore deconvolved the parameter sets in a rather general manner, for which we use a General Mixture model (GMM), a probabilistic model composed of multiple distributions and corresponding weights. We describe each data point as a portion of a normal or log-normal distribution described by

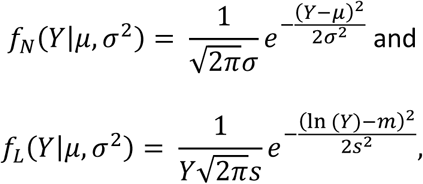

respectively. The logarithmic mean *m* and standard deviation *s* are related to the mean and standard deviation of the normal distribution via [65].

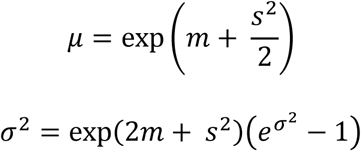

We consider up to three subpopulations to be found in our data and model the total density estimate as a superposition of one, two or three subpopulations, i.e. the Mixture Model reads

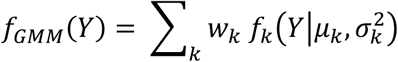

for both normal and log-normal distributions, where to sum goes to 1, 2 or 3, respectively. The variable *w*_*k*_ describes the weights for each population (or component), which satisfy 0 ≤ *w*_*k*_ ≤ 1 and sum up to unity. The weights of each component are directly proportional to the area of the histogram covered by this component and therefore its existence in the data set.

### General Mixture Model analysis

Let *Y* = {*Y*_1_, …, *Y*_*n*_} denote *n* data points. For the scope of this description, assume *Y* to be a one-dimensional variable. Further assume that the data cannot be described by a single distribution, but by a mixture of distributions. A deconvolution of the data into sub-populations faces the following problem: Given a label for each data point, denoting the affiliation to a population, one could group corresponding data points and find the parameters of each population separately using a maximum likelihood estimation or other methods. On the other hand, if we had given the model parameters for each population, labels could in principle be inferred from the likelihood of a data point being described by a population or another. The problem can be formulated by Bayes’ rule (*M* indicates model, *D* indicates data)

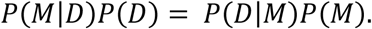

Here, *P*(*M*|*D*) is the posterior probability of the model given the data, which is the aim to calculate. We assign a data point to the component, which maximizes *P*(*M*|*D*). The probability to observe the data given a model is described by *P*(*D*|*M*), i.e. the likelihood function. *P*(*M*) is the prior for the models to be chosen from. In our case, we have no prior beliefs on the models (all models are equally likely) such that *P*(*M*) is uniform. Lastly, the probability *P*(*D*) does not depend on the models and can therefore be dropped.

Neither labels, that is *P*(*M*|*D*), nor model parameters and weights are known a priori. The problem can be approached by an Expectation-Maximization (EM) scheme: Without any prior beliefs about the data distribution, one starts with a simple estimate of model parameters, e.g. a k-means clustering estimate and iterates subsequently between the two following steps until convergence:

#### Expectation step

Calculation of the probability that the component with the current parameter estimate generated the sample, i.e. *P*(*D*|*M*).

#### Maximization step

Update the current parameter estimate for each component by means of a weighted maximum likelihood estimate, where the weight is the probability that the component generated the sample.

We illustrate the results of the EM algorithm exemplary in Additional file 1: Fig S4. From the input data (Additional file 1: Fig S4a), represented as histogram, both the likelihood *P*(*D*|*M*) (Additional file 1: Fig S4b) and the posterior (Additional file 1: Fig S4c) is obtained. The sum of sub-populations corresponds to the overall probability distribution (shown in black) with different model parameters and weights found by maximizing the likelihood function. The posterior describes the probability of data points to fall under each population, i.e. ∑_*k*_ *P*(*M*_*k*_|*D*) = 1. The data points are assigned to those population, for which *P*(*M*_*k*_|*D*) is maximum, resulting in labeled data. The labels are subsequently mapped in two dimensions, visualizing spatial correspondence of slow, intermediate and fast sub-populations (Additional file 1: Fig S4d). The GMM analysis is carried out using the pomegranate machine learning package for probabilistic modeling in Python [66].

#### Selection of subpopulations by the Bayesian Information Criterion (BIC)

A priori, it is not unambiguously clear from how many populations the data is sampled and which form the subpopulations take. We therefore assess the suitability of each model by means of the Bayesian Information Criterion (BIC), which is calculated by [67]

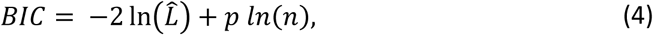

where 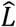 is the maximum likelihood of the maximum likelihood estimation (MLE) estimate, *p* denotes the number of parameters in the model and *n* is the number of data points used for the fit. Among a family of models, the one with the lowest BIC is considered to describe the data best, taking into account competing complexity of models. A large likelihood of a model favors it to describe the data well, while on the other hand the model is penalized if many parameters are involved in the model by the second term in eq. (4). Therefore, the BIC prevents overfitting. In order to judge which model is appropriate for our data, we tested all considered models for each histogram and assessed the optimal model by means of the BIC. The fraction of all histograms which described best by one of the six models considered is given in Table 2. Based on the objective judgement of the fit using the BIC, we chose for each parameter the model which best describes the largest fraction of histograms (Table 2, bold cells).

**Table 2:** Fraction of histograms over all parameters best described by one of the six models considered. The highest fraction is shown in bold.

|  | Normal distribution |  |  | Log-normal distribution |  |  |
| --- | --- | --- | --- | --- | --- | --- |
| #populations | 1 | 2 | 3 | 1 | 2 | 3 |
| D | 0.01 | 0.01 | 0.06 | 0.01 | 0.40 | **0.51** |
| $\alpha$ | 0 | 0.05 | **0.68** | 0.02 | 0.03 | 0.22 |

## AVAILABILITY OF DATA AND MATERIALS

Raw microscopy data is made publicly available [68]. The source code of the Hi-D method is available under the GNU General Public License [69].

## ACKNOWLEDGMENTS

We thank Maxime Dahan (Curie Institute) for gifting the U2OS cell line expressing RPB1, Amy Strom (Gary Karpen Lab, UC Berkeley, USA) for providing images of HP1 in NIH3T3 cells, Silvia Kocanova (CBI Toulouse) for assistance with image acquisition, Raphael Mourad (CBI Toulouse) and Genevieve Fourel (ENS Lyon) for fruitful discussions, Alain Kamgoué (CBI Toulouse) for assistance with computer cluster set-up, and Fatima-Ezzahra L’Faqihi (CPTP-INSERM U1043, TRI) for technical assistance with the Flow cytometry experiments. We acknowledge support from the LITC imaging platform, CBI Toulouse.

## ETHICAL APPROVAL

Not applicable.

## AUTHOR CONTRIBUTIONS

H.A.S. conceived the project, performed the experimental work, developed the data processing algorithm and analyzed the data; R.B, developed the data processing algorithm, wrote the code, carried out simulations and analyzed the data; L.R. performed flow cytometry experiment; H.A.S. and K.B. interpreted the results; H.A.S., R.B. and K.B. wrote the manuscript.

## ADDITIONAL FILES

Additional File 1 (AdditionalFile1.pdf 3.63 Mb) contains details on the conversion from flow fields to trajectories, detailed discussion on the distinction between confined and anomalous diffusion and a comparison of Single Particle Tracking and Optical Flow for reconstruction of motion in intermediate to high labelling density scenarios and further discussion on the comparison of Hi-D and iMSD.

## REFERENCES

1. Serizay J, Ahringer J. ScienceDirect Genome organization at different scales: nature, formation and function. Curr Opin Cell Biol. Elsevier Ltd; 2018;52:145–53.

2. Cremer T, Cremer M, Hübner B, Strickfaden H, Smeets D, Popken J, et al. The 4D nucleome: Evidence for a dynamic nuclear landscape based on co-aligned active and inactive nuclear compartments. FEBS Lett. 2015;589:2931–43.

3. Banerjee B, Bhattacharya D, Shivashankar GV. Chromatin Structure Exhibits Spatio-Temporal Heterogeneity within the Cell Nucleus. Biophys J [Internet]. 2006;91:2297–303. Available from: http://linkinghub.elsevier.com/retrieve/pii/S0006349506719452

4. Fraser P, Bickmore W. Nuclear organization of the genome and the potential for gene regulation. Nature [Internet]. 2007;447:413–7. Available from: http://www.nature.com/articles/nature05916

5. Lieberman-Aiden E, van Berkum NL, Williams L, Imakaev M, Ragoczy T, Telling A, et al. Comprehensive Mapping of Long-Range Interactions Reveals Folding Principles of the Human Genome. Science (80-). 2009;326:289–93.

6. Dixon JR, Selvaraj S, Yue F, Kim A, Li Y, Shen Y, et al. Topological domains in mammalian genomes identified by analysis of chromatin interactions. Nature [Internet]. 2012;485:376–80. Available from: http://www.nature.com/articles/nature11082

7. Beagrie RA, Scialdone A, Schueler M, Kraemer DCA, Chotalia M, Xie SQ, et al. Complex multienhancer contacts captured by genome architecture mapping. Nature. 2017;543:519–24.

8. Rao SSP, Huntley MH, Durand NC, Stamenova EK, Bochkov ID, Robinson JT, et al. A 3D map of the human genome at kilobase resolution reveals principles of chromatin looping. Cell. 2014;159:1665–80.

9. Hansen AS, Pustova I, Cattoglio C, Tjian R, Darzacq X. CTCF and cohesin regulate chromatin loop stability with distinct dynamics. Elife. 2017;

10. Chubb JR, Boyle S, Perry P, Bickmore WA. Chromatin motion is constrained by association with nuclear compartments in human cells. Curr Biol. 2002;12:439–45.

11. Chuang CH, Carpenter AE, Fuchsova B, Johnson T, de Lanerolle P, Belmont AS. Long-Range Directional Movement of an Interphase Chromosome Site. Curr Biol. 2006;16:825–31.

12. Chen B, Gilbert LA, Cimini BA, Schnitzbauer J, Zhang W, Li GW, et al. Dynamic imaging of genomic loci in living human cells by an optimized CRISPR/Cas system. Cell. 2013;155:1479–91.

13. Germier T, Kocanova S, Walther N, Bancaud A, Shaban HAHA, Sellou H, et al. Real-Time Imaging of a Single Gene Reveals Transcription-Initiated Local Confinement. Biophys J. 2017;113:1383–94.

14. Levi V, Ruan Q, Plutz M, Belmont AS, Gratton E. Chromatin Dynamics in Interphase Cells Revealed by Tracking in a Two-Photon Excitation Microscope. Biophys J [Internet]. 2005 [cited 2017 May 23];89:4275–85. Available from: http://www.ncbi.nlm.nih.gov/pubmed/16150965

15. Bornfleth H, Edelmann P, Zink D, Cremer T, Cremer C. Quantitative motion analysis of subchromosomal foci in living cells using four-dimensional microscopy. Biophys J [Internet]. 1999;77:2871–86. Available from: http://www.pubmedcentral.nih.gov/articlerender.fcgi?artid=1300559&tool=pmcentrez&rendertype=abstract

16. Nozaki T, Imai R, Tanbo M, Nagashima R, Tamura S, Tani T. Dynamic Organization of Chromatin Domains Revealed by Super-Resolution Live-Cell Imaging. Mol Cell. 2017;10:1–12.

17. Di Pierro M, Potoyan DA, Wolynes PG, Onuchic JN. Anomalous diffusion, spatial coherence, and viscoelasticity from the energy landscape of human chromosomes. Proc Natl Acad Sci. 2018;115:7753–8.

18. Zidovska A, Weitz D a, Mitchison TJ. Micron-scale coherence in interphase chromatin dynamics. Proc Natl Acad Sci U S A [Internet]. 2013;110:15555–60. Available from: http://www.pnas.org/content/110/39/15555.long

19. Shinkai S, Nozaki T, Maeshima K, Togashi Y. Dynamic Nucleosome Movement Provides Structural Information of Topological Chromatin Domains in Living Human Cells. PLoS Comput Biol [Internet]. 2016;12:1–16. Available from: http://dx.doi.org/10.1371/journal.pcbi.1005136

20. Shaban HA, Barth R, Bystricky K. Formation of correlated chromatin domains at nanoscale dynamic resolution during transcription. Nucleic Acids Res. 2018;46.

21. Monnier N, Guo S-M, Mori M, He J, Lénárt P, Bathe M. Bayesian approach to MSD-based analysis of particle motion in live cells. Biophys J. 2012;103:616–26.

22. Sergé A, Bertaux N, Rigneault H, Marguet D, Sergé A, Bertaux N, et al. Dynamic multiple-target tracing to probe spatiotemporal cartography of cell membranes. Nat Methods. 2008;5:687–94.

23. Di Rienzo C, Gratton E, Beltram F, Cardarelli F. Fast spatiotemporal correlation spectroscopy to determine protein lateral diffusion laws in live cell membranes. Proc Natl Acad Sci. 2013;

24. Cho WK, Jayanth N, English BP, Inoue T, Andrews JO, Conway W, et al. RNA Polymerase II cluster dynamics predict mRNA output in living cells. Elife. 2016;5.

25. Nagashima R, Hibino K, Ashwin SSS, Babokhov M, Fujishiro S, Imai R, et al. Single nucleosome imaging reveals loose genome chromatin networks via active RNA polymerase II. J Cell Biol. 2019;

26. Kim KH, Sederstrom JM. Assaying cell cycle status using flow cytometry. Curr Protoc Mol Biol. 2015;

27. Shukron O, Seeber A, Amitai A, Holcman D. Single particle trajectory statistic to reconstruct chromatin organization and dynamics. bioRxiv [Internet]. 2019;559369. Available from: http://biorxiv.org/content/early/2019/03/04/559369.abstract

28. Amitai A, Seeber A, Gasser SM, Holcman D. Visualization of Chromatin Decompaction and Break Site Extrusion as Predicted by Statistical Polymer Modeling of Single-Locus Trajectories. Cell Rep [Internet]. ElsevierCompany.; 2017;18:1200–14. Available from: https://linkinghub.elsevier.com/retrieve/pii/S2211124717300542

29. Steurer B, Janssens RC, Geverts B, Geijer ME, Wienholz F, Theil AF, et al. Live-cell analysis of endogenous GFP-RPB1 uncovers rapid turnover of initiating and promoter-paused RNA Polymerase II. Proc Natl Acad Sci. 2018;

30. Darzacq X, Shav-Tal Y, de Turris V, Brody Y, Shenoy SM, Phair RD, et al. In vivo dynamics of RNA polymerase II transcription. Nat Struct Mol Biol. 2007;14:796–806.

31. Hajjoul H, Mathon J, Ranchon H, Goiffon I, Mozziconacci J, Albert B, et al. High-throughput chromatin motion tracking in living yeast reveals the flexibility of the fiber throughout the genome. Genome Res. 2013;23:1829–38.

32. Ghosh SK, Jost D. How epigenome drives chromatin folding and dynamics, insights from efficient coarse-grained models of chromosomes. Zhong S, editor. PLOS Comput Biol [Internet]. Public Library of Science; 2018 [cited 2018 Jul 16];14:e1006159. Available from: http://dx.plos.org/10.1371/journal.pcbi.1006159

33. Doi M, Edwards SF. The Theory of Polymer Dynamics. Clarendon Press; 1988.

34. Shukron O, Holcman D. Transient chromatin properties revealed by polymer models and stochastic simulations constructed from Chromosomal Capture data. Marti-Renom MA, editor. PLOS Comput Biol [Internet]. 2017;13:e1005469. Available from: http://dx.plos.org/10.1371/journal.pcbi.1005469

35. Wachsmuth M, Knoch TA, Rippe K. Dynamic properties of independent chromatin domains measured by correlation spectroscopy in living cells. Epigenetics and Chromatin. BioMed Central; 2016;9:1–20.

36. Strom AR, Emelyanov A V., Mir M, Fyodorov D V., Darzacq X, Karpen GH. Phase separation drives heterochromatin domain formation. Nature. 2017;

37. Chen Y. New Approaches for Calculating Moran’s Index of Spatial Autocorrelation. PLoS One. 2013;

38. Manley S, Gillette JM, Patterson GH, Shroff H, Hess HF, Betzig E, et al. High-density mapping of single-molecule trajectories with photoactivated localization microscopy. Nat Methods. 2008;5:155–7.

39. Kind J, Pagie L, Ortabozkoyun H, Boyle S, De Vries SS, Janssen H, et al. Single-cell dynamics of genome-nuclear lamina interactions. Cell. 2013;153:178–92.

40. Akhtar W, De Jong J, Pindyurin A V., Pagie L, Meuleman W, De Ridder J, et al. Chromatin position effects assayed by thousands of reporters integrated in parallel. Cell. 2013;154:914–27.

41. Heitz E. Das Heterochromatin der Moose. Jahrbücher für wissenschaftliche Bot. 1928;69:762–818.

42. Solovei I, Wang AS, Thanisch K, Schmidt CS, Krebs S, Zwerger M, et al. LBR and lamin A/C sequentially tether peripheral heterochromatin and inversely regulate differentiation. Cell. 2013;152:584–98.

43. Falk M, Feodorova Y, Naumova N, Imakaev M, Lajoie RB, Leonhardt H, et al. Heterochromatin drives organization of conventional and inverted nuclei Martin. bioRxiv. 2018;1–19.

44. van Steensel B, Belmont AS. Lamina-Associated Domains: Links with Chromosome Architecture, Heterochromatin, and Gene Repression. Cell. 2017;

45. Kind J, Pagie L, De Vries SS, Nahidiazar L, Dey SS, Bienko M, et al. Genome-wide Maps of Nuclear Lamina Interactions in Single Human Cells. Cell. 2015;

46. Haddad N, Jost D, Vaillant C. Perspectives: using polymer modeling to understand the formation and function of nuclear compartments. Chromosom. Res. 2017. p. 35–50.

47. Sewitz SA, Fahmi Z, Aljebali L, Bancroft J, Brustolini OJB, Saad H, et al. Heterogeneous chromatin mobility derived from chromatin states is a determinant of genome organisation in S. cerevisiae. bioRxiv. 2017;106344.

48. Sakaue T, Saito T. Active diffusion of model chromosomal loci driven by athermal noise. Soft Matter. 2017;

49. Osmanović D, Rabin Y. Dynamics of active Rouse chains. Soft Matter. 2017;

50. Giorgetti L, Galupa R, Nora EP, Piolot T, Lam F, Dekker J, et al. Predictive polymer modeling reveals coupled fluctuations in chromosome conformation and transcription. Cell. 2014;157:950–63.

51. Bensaude O. Inhibiting eukaryotic transcription: Which compound to choose? How to evaluate its activity? Transcription. 2011;

52. Sainsbury S, Bernecky C, Cramer P. Structural basis of transcription initiation by RNA polymerase II. Nat. Rev. Mol. Cell Biol. 2015.

53. Kapranov P, Cheng J, Dike S, Nix DA, Duttagupta R, Willingham AT, et al. RNA maps reveal new RNA classes and a possible function for pervasive transcription. Science (80-). 2007;

54. Lu Z, Lin Z. Pervasive and dynamic transcription initiation in Saccharomyces cerevisiae. Genome Res. 2019;

55. Cisse II, Izeddin I, Causse SZ, Boudarene L, Senecal A, Muresan L, et al. Real-time dynamics of RNA polymerase II clustering in live human cells. Science (80-). 2013;341:664–7.

56. Lukinavičius G, Blaukopf C, Pershagen E, Schena A, Derivery E, Gonzalez-gaitan M, et al. A far-red DNA stain for live-cell nanoscopy. Nat Commun. 2015;61:3–5.

57. Eddaoudi A, Canning SL, Kato I. Flow cytometric detection of g0 in live cells by Hoechst 33342 and Pyronin Y staining. Methods Mol Biol. Humana Press Inc.; 2018. p. 49–57.

58. Tomasi C, Manduchi R. Bilateral Filtering for Gray and Color Images. Proc 1998 IEEE Int Conf Comput Vis. 1998;

59. Saxton MJ, Jacobson K. Single-particle tracking: applications to membrane dynamics. Annu Rev Biophys Biomol Struct. 1997;26:373–99.

60. Dickerson D, Gierlinski M, Singh V, Kitamura E, Ball G, Tanaka TU, et al. High resolution imaging reveals heterogeneity in chromatin states between cells that is not inherited through cell division. BMC Cell Biol [Internet]. 2016;17:33. Available from: http://bmccellbiol.biomedcentral.com/articles/10.1186/s12860-016-0111-y

61. Seber GAF, Wild CJ. Nonlinear Regression [Internet]. John Wiley Sons. 2003. Available from: http://books.google.com/books?id=YBYlCpBNo_cC&pgis=1

62. He J, Guo S-M, Bathe M. Bayesian approach to the analysis of fluorescence correlation spectroscopy data I: theory. Anal Chem [Internet]. 2012;84:3871–9. Available from: http://pubs.acs.org/doi/abs/10.1021/ac2034369%5Cnpapers2://publication/doi/10.1021/ac2034369%5Cnhttp://www.ncbi.nlm.nih.gov/pubmed/22423978

63. Rousseeuw PJ, Croux C. Alternatives to the Median Absolute Deviation. J Am Stat Assoc [Internet]. 1993 [cited 2017 Apr 25];88:1273–83. Available from: http://www.tandfonline.com/doi/abs/10.1080/01621459.1993.10476408

64. Vrljic M, Nishimura SY, Brasselet S, Moerner WE, McConnell HM. Translational diffusion of individual class II MHC membrane proteins in cells. Biophys J [Internet]. 2002;83:2681–92. Available from: http://www.ncbi.nlm.nih.gov/entrez/query.fcgi?cmd=Retrieve&db=PubMed&dopt=Citation&list_uids=12414700

65. Mood AM, Graybill FA, Boes DC. Introduction to the Theory of Statistics. Book [Internet]. 1974;3:540–1. Available from: http://www.librarything.com/work/1154157/book/32217714

66. Schreiber J. Pomegranate: fast and flexible probabilistic modeling in python. 2017 [cited 2018 Jun 27]; Available from: http://arxiv.org/abs/1711.00137

67. Schwarz G. Estimating the Dimension of a Model. Ann Stat [Internet]. Institute of Mathematical Statistics; 1978 [cited 2018 May 6];6:461–4. Available from: http://projecteuclid.org/euclid.aos/1176344136

68. Shaban HA, Barth R, Recoules L, Bystricky K. Hi-D: Nanoscale mapping of nuclear dynamics in single living cells. Datasets. https://doi.org/10.6084/m9.figshare.11793801.v1

69. Shaban HA, Barth R, Recoules L, Bystricky K. Hi-D: Nanoscale mapping of nuclear dynamics in single living cells. Github. doi: 10.5281/ZENODO.3634347 (2019)

