## Supplementary file for "Hi-D: Nanoscale mapping of nuclear dynamics in single living cells"

### ADDITIONAL FILE 1

### CONTENTS

**NOTE S1** P. 4

This note provides detailed information about the conversion of flow fields into trajectory.

**NOTE S2** P. 8

This note provides a detailed view on the comparison of confined and anomalous diffusion, which is supported by exemplary simulations of the two types of diffusion.

**NOTE S3** P. 13

This note provides a detailed information about simulations carried out to compare Optical Flow and Single Particle Tracking reconstruction of trajectories in different imaging conditions and illustrate the response of Optical Flow to experimental issues such as diffusion out of focus and heterogeneous labelling density in the field of view.

**NOTE S4** P. 23

This note provides an explanation and an intuitive illustration employing a simulated chromosome of why discrepancies between Hi-D and SPT approaches can usually arise.

**NOTE S5** P. 25

This note provides more information and a detailed discussion about the comparison of Hi-D to iMSD.

**SUPPLEMENTARY REFERENCES** P. 34

#### SUPPLEMENTARY FIGURES & MOVIES

**Fig S1:** Conversion from flow fields to trajectories. P. 6

**Fig S2:** Illustration of MSD and Bayesian selection illustration for parameter mapping. P. 7

**Fig S3:** Comparison of confined and anomalous diffusion. P. 10

**Fig S4:** General Mixture Model analysis. P. 11

**Fig S5:** Simulation workflow and exemplary evaluation. P. 18

**Fig S6:** Size distribution of heterochromatin regions. P. 19

**Fig S7:** Comparison of Optical Flow and Single Particle Tracking by simulated image series. P. 21

**Fig S8**: The effect of varying signal-to-noise ratios on the estimation of dynamic parameters with the Hi-D approach. P.26

**Fig S9:** The effect of varying time intervals between frames on the estimation of dynamic parameters with the Hi-D approach. P.27

**Fig S10:** Addition of serum increases transcription without affecting drastically cell cycle phases. P. 29

**Fig S11:** Chromatin diffusion is reduced at the nuclear periphery compared to the nuclear interior. P. 30

**Fig S12:** Hi-D analysis for a different cell line (MCF7 cells). P. 31

**Fig S13:** Dynamics of H2A and HP1α in NIH3T3 cells confirm that chromatin dynamics is independent from compaction, but suggest that eu- heterochromatin phase-separation forms a barrier for chromatin mobility. P. 32

**Movie S1**: Illustration of discrepancies due to different imaging modalities between SPT methods and Hi-D using a simulated chromosome. P. 24

#### NOTE S1

##### Conversion of flow fields into trajectories

In high-density scenarios where Single Particle Tracking methods reach their limit, dense Optical Flow methods present a powerful tool to investigate local bulk motion of biological macromolecules. Here we determine flow fields of fluorescently labelled DNA and reconstruct virtual trajectories to extract motion at sub-pixel resolution and long-time intervals at the level of the whole nucleus [1]. Optical Flow algorithms estimate motion between frames as a field description (Eulerian description) of the underlying continuum motion, evaluated at fixed 'stations', i.e. the pixel positions in the Cartesian coordinate system, which is a powerful approach when the coordinates of single particles cannot be defined. In contrast, actual tracking of particles' coordinates over time is referred to as Lagrangian description. A continuum motion consisting of a finite number of particles can be described in both ways, according to continuum mechanics [2–4]. Because it is impossible to identify individual emitters (or particles) in densely labelled images, we start out with a limited number of *virtual* particles, which are assumed to be seeded on a regular grid which is defined by the pixels. Let $\vec{r}_0$ denote these (fixed) pixel positions. Let the coordinates of the particle in Cartesian space be $\vec{\xi}_{r_0}(t)$ at any time point $t$. Consider further that the Eulerian flow field is known only at the positions $\vec{r}_0$, but can be evaluated at any position by interpolation of the coordinates of interest. Then, the particle's (Lagrangian) velocity $\vec{q}_L(\vec{\xi}_{r_0}, t)$ at position $\vec{\xi}_{r_0}$ and time $t$ is the same as the Eulerian velocity at $\vec{\xi}_{r_0}(t)$

$$\vec{q}_L(\vec{\xi}_{r_0}, t) = \frac{\partial \vec{\xi}_{r_0}}{\partial t} \tag{1}$$

Therefore, the trajectory, consisting of the consecutive positions of the particle can be obtained by integration of Equation (1) using the fact that Eulerian and Lagrangian velocities are equal when evaluated at the same position. In situations where particle detection is impossible (e.g. due to high density of emitters), the Eulerian description of continuum motion can be translated to a Lagrangian description by considering *virtual* particles and using the flow field description to extract their hypothetical trajectories. We consider virtual particles with initial positions at the center of each image pixel, for which flow was estimated by Optical Flow (Fig S1a, first flow field highlighted). Note that the flow fields describe the motion *between* frames, whereas particle coordinates are described at the imaging time of each frame. The time evolution, i.e. the trajectory of each virtual particle is reconstructed as follows: From the particle's initial position $\vec{\xi}_{r_0}(t_0) = \vec{r}_0$, the flow field dictates the displacement from frame 1 to frame 2 (dark blue trajectory segment in Fig S1d, e), i.e. $\vec{\xi}_{r_0}(t_1) = \vec{r}_0 + \vec{q}_L(\vec{r}_0, t_0)\,\Delta t$, where $\Delta t = t_{i+1} - t_i$

denotes the time between consecutive frames. The current particle coordinates at $t_1$ do not necessarily coincidence with the regular grid on which the flow field is evaluated. We therefore interpolate the flow field at time $t_1$ to the particle coordinates $\vec{\xi}_{r_0}(t_1)$ (Fig S1b), light blue flow field) and can evaluate the particle coordinates at $t_2$: $\vec{\xi}_{r_0}(t_2) = \vec{\xi}_{r_0}(t_1) + \vec{q}_L\big(\vec{\xi}_{r_0}(t_1), t_1\big)\,\Delta t$, where $\vec{q}_L\big(\vec{\xi}_{r_0}(t_1), t_1\big)$ denotes the interpolated flow field (i.e. displacement vectors) at time $t_1$ at the particle coordinates. This procedure is repeated until all flow fields are processed (Fig S1c) and the resulting visual particle coordinates are connected to form trajectories (Fig S1d, e). Note that extrapolation outside the nucleus and in nucleoli where no signal intensity and therefore no flow field is given, is not considered.

Importantly, the concept of virtual particles does not correspond to particles associated with specific foci along the genome. These are impossible to detect with the given labelling density. Instead, virtual particles move along a trajectory which reflects the local bulk motion of many emitters as computed by Optical Flow. Indeed, it is highly likely that a single pixel may contain fluorescence signal from more than one chromatin fibre and thus calculated trajectories should not be interpreted in the sense of single particle tracking trajectories. Instead, trajectories across multiple pixels in a local neighbourhood have to be taken into account in order to interpret quantities derived from these trajectories.

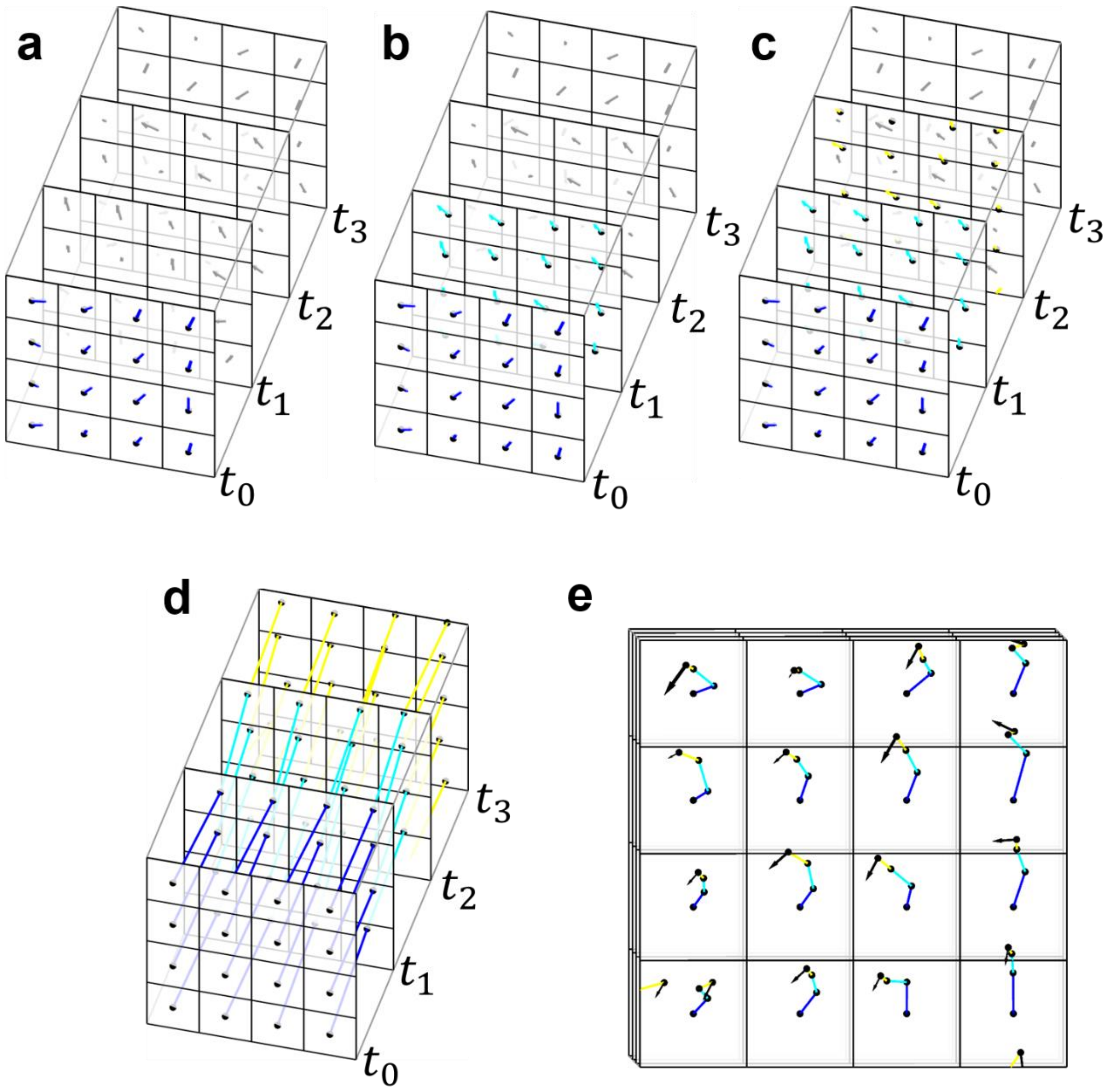


**Fig S1: Conversion from flow fields to trajectories. a)** Flow fields are evaluated on a fixed grid given by the pixels of the input images to Optical Flow. The first flow field of the series is highlighted in dark blue. Virtual particles whose initial coordinates coincidence with the center of image pixels are displaced by the flow field at $t_0$. **b)** The flow field at $t_1$ is interpolated on the current positions of the virtual particles and displaced according to the interpolated flow field. **c)** The procedure is repeated for all flow fields in the series resulting in subsequent positions of virtual particles given by the displacement of flow fields. **d)** Particle positions are reconnected to form trajectories. The color of trajectory segments denote the flow field which was used to displace the particles from $t_i$ to $t_{i+1}$. **e)** Quasi-2D representation of virtual particle trajectories over time as shown in d).

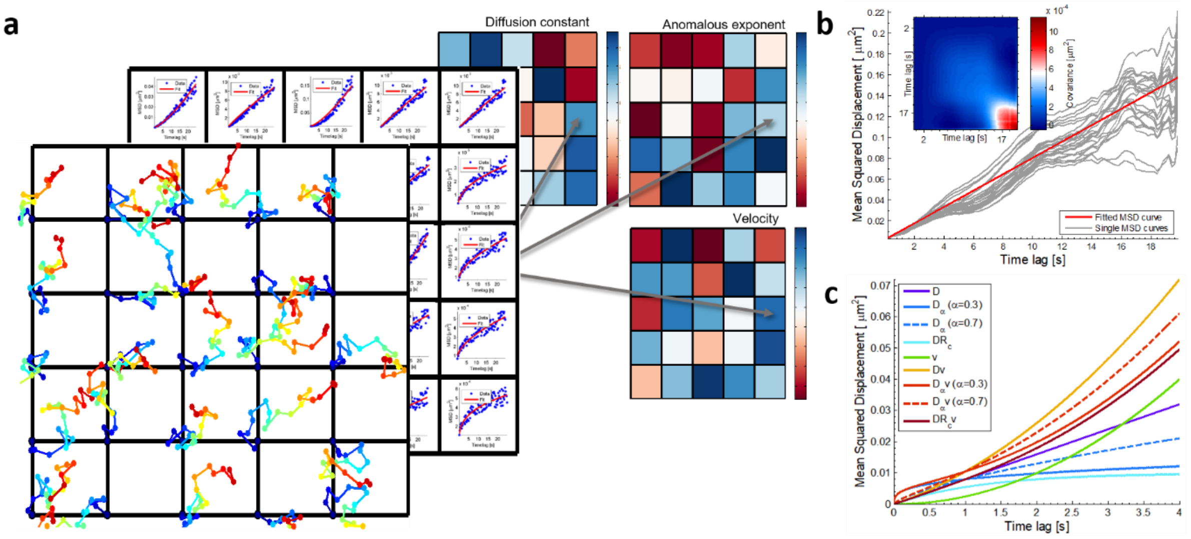


**Fig S2: Illustration of MSD and Bayesian selection illustration for parameter mapping. a)** The trajectory of a virtual particle in every pixel inside the nucleus is calculated by using previously determined flow fields. For every pixel, the MSD is calculated. Fitting the MSD, taking into account its neighbourhood, results in a set of parameters (diffusion coefficient, anomalous exponent, drift velocity) for each pixel. **b)** The MSD curves for one example pixel and its neighbouring pixels. Because errors are correlated in MSD calculation, the covariance matrix is calculated for the adjacent MSD curves (inset). The Cholesky decomposition of the covariance matrix is used in a generalized least squares approach to transform the MSD model and experimental data into a coordinate system, in which the errors are no longer correlated and an ordinary least squares fit is appropriate to find a solution to the optimization problem. Note that the MSD calculation for large time lags has less statistical power than for small time lags due to the lack of pairs to average. Therefore, the covariance matrix shows a high variation for large time lags in turn resulting in small weights for large time lags. The fitted MSD curve is shown in red. **c)** Models to describe the empirical MSD for some exemplary parameter values.

#### NOTE S2

##### Comparison of confined and anomalous diffusion

In order to test the ability of the Bayesian classification routine used in this study to resolve confined and anomalous diffusion (DR and DA respectively), we simulate particles undergoing either one of the two types of diffusion. Confined diffusion is characterized by a particle diffusing freely within a sphere of radius $R_c$ with diffusion constant $D$. The space outside the sphere has the form of an infinite potential impossible to overcome and hence resulting in confinement to the volume of the sphere. Exemplary simulated trajectories are shown in two dimensions in Fig S3a-d (left column) for different values of diffusion constant and radius of confinement. Anomalous diffusion is characterized by an effective potential exerting a driving force towards the particle's origin, whose source may be to surround obstacles hindering free diffusion. We model the driving potential as a harmonic potential with the characteristic dimension $L_{trap}$. The particle feels a spring-like driving force with spring constant $\propto L_{trap}^{-2}$. Exemplary trajectories for anomalous diffusion are shown in Fig S3a-d (middle column) for different values of $L_{trap}$. The potential strength is indicated by colour code. The theoretical MSD is given as in equations 1 and 3 of the Methods section for anomalous and confined diffusion respectively.

For confined diffusion, one can approximate the expression for short and long time scales:

$\boldsymbol{D\tau \ll R_c^2}$**:** The exponential can be expanded in a Taylor series yielding $MSD_{DR}(\tau) \approx R_c^2\left(1-\left(1-\frac{4D\tau}{R_c^2}+\cdots\right)\right) \approx 4D\tau$, i.e. free diffusion in first order. Effectively, the particle does not feel any confinement for short time lags as the explored volume is much smaller than the confinement volume.

$\boldsymbol{D\tau \gg R_c^2}$**:** The exponential argument is small and therefore $MSD_{DR}(\tau) \approx R_c^2(1-1) = R_c^2$. The confinement is effectively a hard wall potential, impossible to overcome for the particle. For long time lags, the particle therefore explored the whole available volume, but cannot reach any further, resulting in a constant MSD (see Fig S2c).

For anomalous diffusion, the particles move freely for short times, but adapt sub-diffusive behaviour for longer times as the effect of the external potential becomes dominant. However, a particle undergoing anomalous diffusion is not confined in the sense of a hard wall potential and can in principle diffuse in all space [5]. This leads to a continuously rising MSD for $\tau \to \infty$. Despite the analytical form of the MSD for confined and anomalous diffusion (exponential versus algebraic), the behaviour for long time lags is a main characteristic for the distinction of the two types of diffusion.

In order to illustrate the theoretical curve shape, exemplary scenarios are shown in Fig S3a-d, where different levels of mobility and confinement / anomaly and the ability to resolve the correct type of motion by means of the MSD (right column) are explored. For the scenarios a-c) the limit $D\tau \ll R_c^2$ is not reached within a trajectory length of 150 steps and the shapes of mean MSD for confined (green) and anomalous diffusion (red) are similar. Consequently, the exact type of diffusion could not be resolved. However, for strong confinement (Fig S3d), the curves are sufficiently dissimilar and allow extracting the correct MSD model. Experimentally, only a finite trajectory length can be recorded and it is questionable when the trajectory length is sufficient to observe the long time lag limit when no prior information about the particle environment is known. To test whether a reliable distinction can be made for experimentally observed trajectories, an example nucleus is analysed twice. First, the Bayesian classifier is given the choice between free, directed and confined diffusion (and combinations thereof: D, DR, V, DV, DRV). The model selection is shown in Fig S3e (left). The majority of trajectories are classified as confined + directed diffusion (DRV), whereas only about 25% of trajectories are classified as purely confined diffusion. Next, the model for confined diffusion is replaced by anomalous diffusion and the analysis is carried out again. First of all, high agreement between the two modes of analysis is seen for trajectories classified as purely Brownian and directed as well as a combination thereof. However, the fraction of anomalous + directed diffusion (DAV) is small compared to purely anomalous diffusion. In particular, about 92% of trajectories classified as DAV were previously classified as DRV. From the remaining trajectories not classified DAV, about 72% are classified as purely anomalous diffusion. These results suggest that only a subset of trajectories classified as combination of confined and directed diffusion is consistent with the combination of anomalous and directed diffusion. The majority of trajectories classified as DRV is preferentially described by purely anomalous diffusion. A reason might be that in the case where only confined diffusion is allowed to describe experimental trajectories, an effective directed transport is needed to account for the continuous rise of the MSD even for large time lags. Finally, it remains unclear if confined or anomalous diffusion is present in experimental trajectories as long as the plateau in the MSD is not reached. Even though strong experimental evidence for confined diffusion of proteins and molecules exists [6] for large biomolecules such as chromatin, a confinement may have several forms such as anisotropic or temporally varying confinement radii with to date unknown sources. The idealized model of a hard wall potential defining the confinement volume may not be appropriate for most biological cases (except for membranes) resulting in a mixed state between confined and anomalous diffusion, which is hard or even impossible to resolve without high spatiotemporal resolution and long-time measurements. The exemplary results and reasoning above suggest that sub-diffusive behaviour observed for chromatin may be best described by anomalous diffusion rather than confined diffusion to prevent misclassification and misinterpretation.

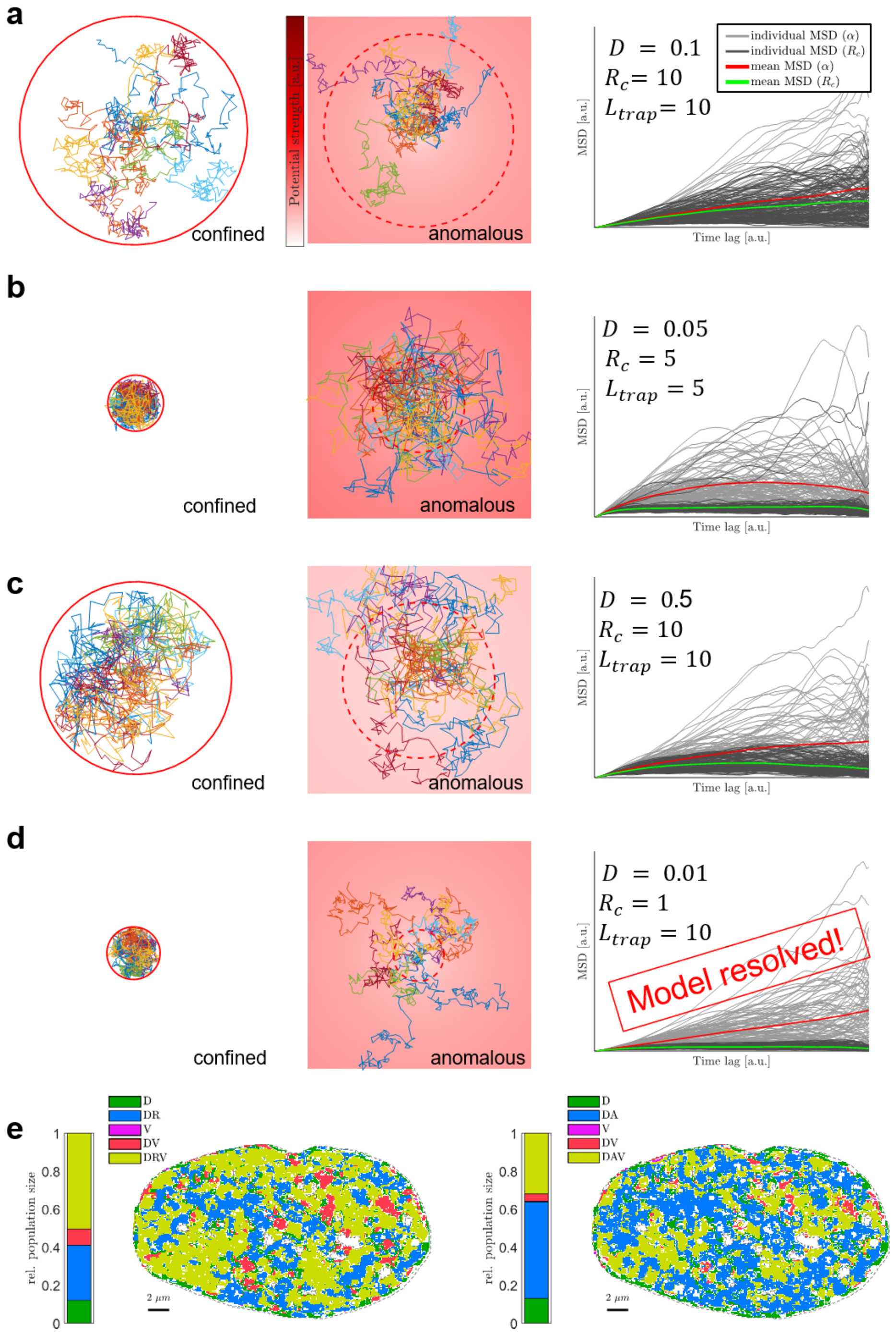


**Fig S3: Comparison of confined and anomalous diffusion.** Confined (DR) and anomalous diffusion (DA) was simulated in three dimension for 100 particles over 150 frames for different degrees of confinement / anomaly. The resulting MSD curves for anomalous and confined motion are calculated and were input to the Bayesian model classifier used in this study, where the model choice was only left between pure anomalous and confined motion. 10 example trajectories are shown as projection to two dimensions for confined motion (left column) and anomalous motion (middle column). The confinement radius is indicated by a circle (left column) and for comparison as dashed circle (middle column), respectively. The back-driving potential is indicated by color code. Individual and average MSD curves are shown for all trajectories. The following parameter pairs are shown: **a)** moderate diffusion and weak confinement / anomaly, **b)** slow diffusion and moderate confinement / anomaly, **c)** slow diffusion and weak confinement, **d)** very slow diffusion and strong confinement, but weak anomaly. All numerical values in a.u. **e)** An example nucleus stained with DNA-Sir analyzed in two modes: The Bayesian model selection is allowed to choose from free, directed, confined diffusion and corresponding combinations (left) or anomalous diffusion (right) as indicated by color and summarized as stacked bars.

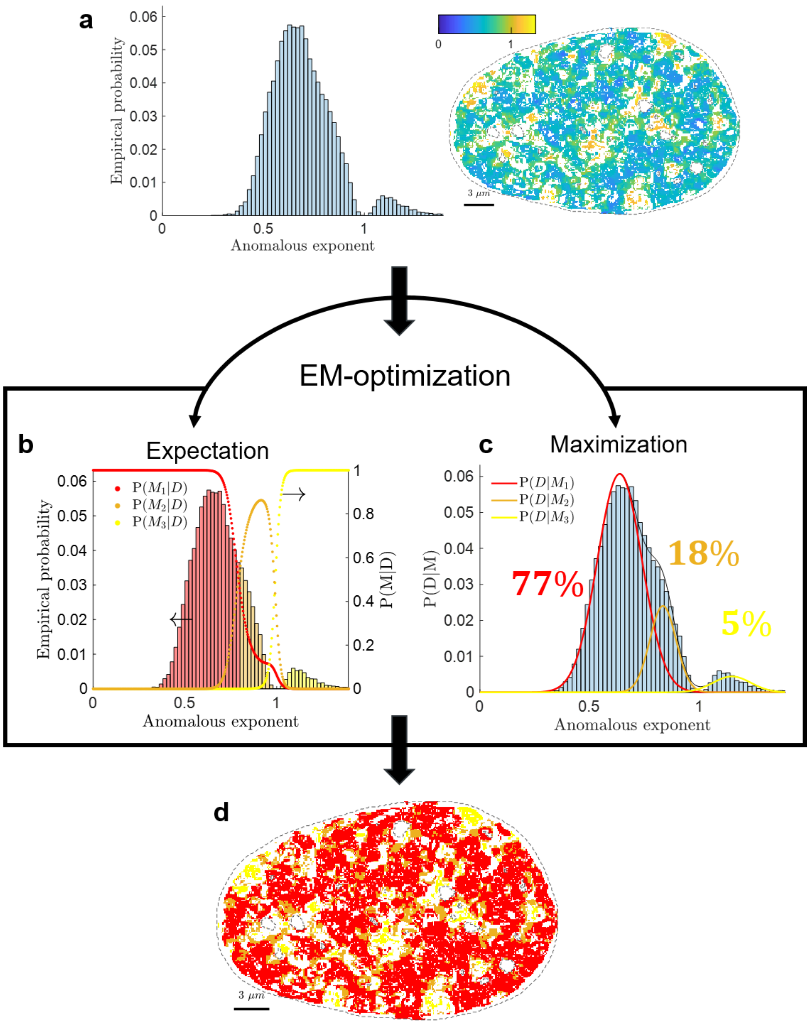


**Fig S4**: General Mixture Model analysis. **a)** An example nucleus showing the spatial distribution of the anomalous exponent across the nucleus (right) and the corresponding histogram of the data with its empirical probability. The General Mixture Model analysis aims at estimating the parameters of underlying populations within the data, here given as three independent Gaussian distributions. Initial estimates are computed by a k-means clustering. The Expectation-Maximization (EM) algorithm iteratively refines data labels and population parameters to maximize the marginal likelihood of observed data until convergence. **b)** Calculating the probability of each data point to belong to one of the populations given the current parameter estimates of each population. The probability sums up to one for each data point and is shown for the three populations on the right axis. **c)** Given a labeling of data into populations, refine the population parameters by weighted maximum likelihood estimates. **d)** Each data point is labeled according to its maximal probability to belong to one of the populations, i.e. $\mathrm{M} = \mathrm{argmax_k} P(M_k|D)$ and is mapped back to the two-dimensional spatial distribution.

#### NOTE S3

##### Performance comparison of Single Particle Tracking and Optical Flow by simulation

In order to quantitatively evaluate the performance of Optical Flow (OF) and Single Particle Tracking (SPT) methods with respect to trajectory reconstruction, ground truth (GT) data is necessary, which is in general not available in experimental data. For this purpose, we simulated typical fluorescence microscopy images of particles undergoing realistic motion in three dimensions (Fig S5). We consider three scenarios: low and high density of emitters as well as a high density scenario with local patches of super-high density (Fig S5, Fig S7). After simulation of an image series, the complete series is input to the two trajectory reconstruction algorithms. In order to compare the dynamic properties of all considered trajectories, the MSD is calculated and the following three quantities are computed: Cosine similarity, relative Euclidian distance and the relative error of diffusion constant, which was derived from a regression of the MSD. These measures allow for a comparison of the MSD shape, a characteristic of the underlying type of motion revealed by the MSD, as well as systematic over- and underestimation of local dynamics. In the following, we describe the simulation procedure, evaluation measures which were used to assess the accuracy of both methods as well as the results.

##### Simulation of microscopy images

**Emitter density** The emitter density has a limiting influence on the performance of SPT methods, as shown in an extensive comparison study of SPT methods [7]. We consider two density levels: $0.001/px^3$ (low density) and $0.02/px^3$ (high density). In order to further mimic high heterogeneity of chromatin density, we simulate local regions of even higher density ($0.035/px^3$), corresponding to eu- and heterochromatin domains. The density of heterochromatin was chosen such as to match a nucleosome density ratio of 0.58 determined previously [8]. The proportion of heterochromatin domains was empirically determined using a volume proportion of eu- and heterochromatin of about 12.5 % [8]. This enables determining regions of heterochromatin within a fluorescence microscopy image as described previously by Wachsmuth *et al.* [8]. In brief, the image is blurred by a Gaussian Filter and the 12.5% of the highest intensity value is extracted indicating high chromatin density within these pixels. Fig S6 shows an example nucleus with regions of heterochromatin as determined by Wachsmuth *et al.;* the heterochromatin regions are marked in white and individual areas are found (Fig S6b, c), from which the area of each domain is calculated for all nuclei used in this study. The area distribution was found to be well described by an exponential distribution with mean $1.9 \pm 3.5\,\mu m^2$ (Fig S6d). This value was a guideline for the simulation of artificial heterochromatin domains, which were assumed to be circular for simplicity. The number of areas was adjusted such that the total area of heterochromatin domains corresponds to about $12.5\%$ of the total area of the simulated volume (as projected to two dimensions).

A volume of 128 x 128 x 15 pixels is simulated corresponding to approximately 8 x 8 x 1 $\mu m$ (pixel size $\sim 65\ nm$). Typically, the depth of the focal plane in a confocal microscopy image is about $\sim 500\ nm$; we therefore simulate twice this thickness in order to allow emitters to move in and out of the focal plane. The mentioned parameters result in an average number of particles of about 240, 4770 and 5500 particles respectively for the three scenarios, which are seeded randomly in the simulated volume. Particles are not seeded on the image boundary to avoid boundary effects. An initial example configuration for the scenario of super-high density patches in shown in Fig S5a.

**Particle dynamics** We simulate Brownian motion in three dimensions such that particles are allowed to randomly move in and out of the imaged volume. A diffusion coefficient of $D = 5 \cdot 10^{-3}\ \mu m^2/s$ was used, matching previously determined values for the diffusion constant for chromatin [9,10]. A total of 20 frames were simulated governing $4\ s$ of experimental imaging (acquisition time $\tau = 200\ ms$). Therefore, particle displacements between subsequent images were drawn from a normal distribution with mean zero and variance $\sigma^2 = \sqrt{6D\tau} \approx 1.2\ px$. It was previously shown [1] that chromatin motion shows a high degree of correlated motion, which allows to empirically impose motion correlation to the purely random Brownian dynamics. We therefore displace particles within a domain of correlated motion collectively, i.e. several emitters along the same displacement vector for each frame independently as reported previously [1]. Furthermore, particle appearance and disappearance are regulated by random processes [7] due to several factors such as emitter transitions into a dark triplet state. Seeding N particles uniformly over the simulation volume, particle disappearance was modeled by a Bernoulli process with probability $\alpha = 0.05$ for every particle to disappear at each time step and the possibility of reappearance excluded. On the other hand, particles may appear at random locations at every time step and the number of appearing particles is drawn from a Poisson distribution with mean $N\alpha$, such that the number of appearing and disappearing particles balance each other. From all generated trajectories, those with a length of 4 or less frames were discarded.

**Imaging process** Due to a diffraction limited optical microscopy setup, the imaging of fluorescent photons is modelled as the convolution of the light emission field and the point spread function (PSF) of a typical confocal microscope. The PSF of an optical system is the image of a point source and the pupil function is defined as the Fourier transform thereof. Therefore, the complex-valued amplitude point spread function $PSF_A$ and the pupil function $P$ form a Fourier transform pair:

$$P(k_x, k_y) = \iint_{-\infty}^{\infty} PSF_A(x, y) \cdot e^{-2\pi i(k_x x + k_y y)} dx dy$$
$$PSF_A(x, y) = \iint_{-\infty}^{\infty} P(k_x, k_y) \cdot e^{2\pi i(k_x x + k_y y)} dk_x dk_y \tag{2}$$

In other words, the PSF can easily be computed by the use of discrete Fourier transforms if the pupil function of the optical system is known. The complex pupil function in our case is simply described by a disk with a radius, which is defined by the ratio of numerical aperture NA and the wavelength of the light $\lambda$:

$$P(k_x, k_y) = e^{2\pi i} \cdot \begin{cases} 1 & for\ k_x^2 + k_y^2 \leq \frac{2\pi NA}{\lambda} \\ 0 & otherwise \end{cases}$$

The treatment above is carried out in scalar diffraction theory and is idealized assuming equal excitation and emission wavelength, no aberration and a constant value inside the pupil function disk. Furthermore, the point source is assumed to be in focus. Emission outside the focal plane is included by adding a defocus to the PSF. This is done by separately expressing $k_z(k_x, k_y) = \sqrt{\left(\frac{n}{\lambda}\right)^2 - k_x^2 - k_y^2}$ , where $n$ is the refractive index of the immersion medium and multiplying the integrand in eq. (2) by a 'defocus phase function' $\exp(2\pi i k_z(k_x, k_y) z)$ such that the expression for the amplitude point spread function in three dimensions reads [11]

$$PSF_A(x, y, z) = \iint_{-\infty}^{\infty} P(k_x, k_y) \cdot e^{2\pi i(k_x x + k_y y)} e^{2\pi i k_z(k_x, k_y) z} dk_x dk_y$$

The observed PSF is the absolute square of the computed amplitude PSF, i.e. $PSF = |PSF_A|^2$. The parameters defining the shape of the PSF are set as follows: emission wavelength $\lambda = 647\ nm$, numerical aperture $NA = 1.4$, refractive index of immersion medium $n = 1.3$ and pixel size $65\ nm$. The simulated PSF is shown in (Fig S5b) for exemplary z-slices, and a convolved simulation volume is shown in (Fig S5c).

Next to blurring, the imaging process is subject to unavoidable noise. In practice, two predominant sources of noise exist, namely signal-dependent Poisson noise and setup-dependent Gaussian white noise due to several factors such as camera gain and thermal noise. The signal-to-noise ratio (SNR) determines the presence of noise photons in contrast to signal photons and is defined as the ratio of squared signal intensity $I^2$ and noise variance $\sigma^2$ on a decimal logarithmic scale: $SNR = 10 \log_{10}(I^2/\sigma^2)$ . While Poisson noise is dependent on the number of observed photons, the Gaussian noise contribution can be varied to match the SNR of about $21\ dB$, which we typically observe in our data. The final image is the projection of the whole convolved simulation volume in two dimensions with applied Poisson and Gaussian noise (Fig S5d).

**Performance measures**

**Cosine Similarity** The cosine similarity addresses the orientation similarity between two multidimensional vectors, or in our case between two MSD curves. Defining the two MSD curves to compare as arrays $\vec{a}$ and $\vec{b}$, the cosine similarity is defined as

$$\cos\theta = \frac{\vec{a} \cdot \vec{b}}{||\vec{a}||\ ||\vec{b}||}.$$

The cosine similarity returns 1 if the two curves have the same shape, regardless of their magnitude and values < 1 otherwise. The shape of the MSD curve has an important meaning as it is characteristic for the underlying type of motion. For instance, motion is considered Brownian if the MSD follows a linear relationship over time lag, whereas a deviation denotes constraint (sub-linear relationship) or active transport (super-linear). In case that multiple estimated trajectories are associated to a single GT trajectory, the mean MSD curve is compared to GT.

**Relative Euclidean distance** The Euclidean distance between two multidimensional vectors is defined as the ratio of the Euclidean distance between the vectors and the norm of the GT to compare to. Denoting the GT by $\vec{a}$ and the estimated MSD by $\vec{b}$:

$$ED = \frac{||\vec{a} - \vec{b}||}{||\vec{a}||}$$

The Euclidean distance is measure of magnitude yielding 0 if the GT and estimated MSD coincide perfectly. Otherwise, the relative Euclidean distance returns the norm of the deviation between the two MSD curves with respect to the GT. In case that multiple estimated trajectories are associated to a single GT trajectory, the mean MSD curve is compared to GT.

**Estimated relative diffusion coefficient** Another obvious measure to assess the accuracy of the reconstruction of particle motion is the diffusion coefficient of GT and estimated trajectories. Diffusion is a stochastic process, such that in general a single trajectory cannot display the average simulated behavior of many particles. Both SPT and OF determine the local motion of particles rather than an ensemble average. For this reason, it is appropriate to compare local quantities rather than comparing to an average value. We therefore face the problem of fitting a single GT MSD and possibly several estimated MSD curves ultimately comparing the diffusion constant derived from the regressions. In case that only 1 MSD curve is found to be fit, which is the case for the GT curve and most of the estimated SPT trajectories, a weighted nonlinear regression is used, weights being the standard deviation of the MSD curves. For more than 1 but less than 5 trajectories, the mean MSD is used for fitting. In case that more than 5 trajectories have been found, an appropriate way of fitting is described in (see main text, Methods). We consider only the free diffusion model in order to be able to compare the diffusion constant to the weighted non-linear regression. Similar to the relative Euclidean distance, we compute the relative error between estimated and simulated MSD curves after regression as

$$\sigma_{\mathrm{D}} = \frac{D - D^*}{D^*},$$

where $D^*$ is the GT and $D$ is the reconstructed diffusion coefficient. By the regression, further inaccuracy is introduced, but a comparison is nevertheless possible and an overall trend is apparent. To see how these performance measures, reflect the behaviour of individual trajectories and corresponding MSD curves, three example trajectories reflecting different cases of the motion reconstruction by OF and SPT compared to the GT trajectory are shown in Fig S5e. The performance measures defined above are given in **Fehler! Verweisquelle konnte nicht gefunden werden.**1 for the three example trajectories. The direction of OF and SPT are similar for approximately the first half of the trajectory (panel i), whereas the GT trajectory propagates in the opposite direction. For longer time points the OF trajectory resembles the one of the GT, whereas the SPT trajectory deviates largely, possibly due to a reconnection error. The MSD curve reflects the similarity of GT and OF estimation despite of the deviation in direction of the trajectories' propagation leading to relatively accurate performance measures. The reconnection error in SPT leads to a tremendous increase in MSD values especially for long time lags yielding a considerably lower cosine similarity and more than 400% error in Euclidean distance and estimation of diffusion constant. Disappearance of a GT emitter after 7 frames in shown in panel (ii). Both OF and SPT detect trajectories throughout the complete frame series using intensity information from surrounding emitters. Motion is accurately estimated by OF in the first 5 time lags and largely deviates from the GT MSD until its disappearance. In this case, the shape of the MSD is well reflected by the SPT trajectory leading to a considerably higher value in cosine similarity for SPT than for OF. However, SPT overestimated the motion leading to large errors in the relative Euclidean distance and diffusion coefficient, whereas the resembling of GT and OF in the first few time lags leads to a relatively accurate estimate for the diffusion coefficient due to a small standard deviation and therefore large weight of the OF MSD curves for small time lags. An example of a false negative of SPT and therefore no dynamic information for the SPT data set is shown in panel (iii).

| | (i) | | (ii) | | (iii) | |
|---|---|---|---|---|---|---|
| | OF | SPT | OF | SPT | OF | SPT |
| Cosine similarity | 0.991 | 0.898 | 0.877 | 0.956 | 0.997 | - |

| | | | | | | |
|---|---|---|---|---|---|---|
| Rel. Euclidean distance | 0.31 | 4.62 | 0.88 | 5.28 | 0.09 | - |
| Rel. error in D | -0.72 | 4.57 | 0.13 | 5.44 | -0.68 | - |

**Table S1:** Performance measures for example trajectories shown in Fig S5e

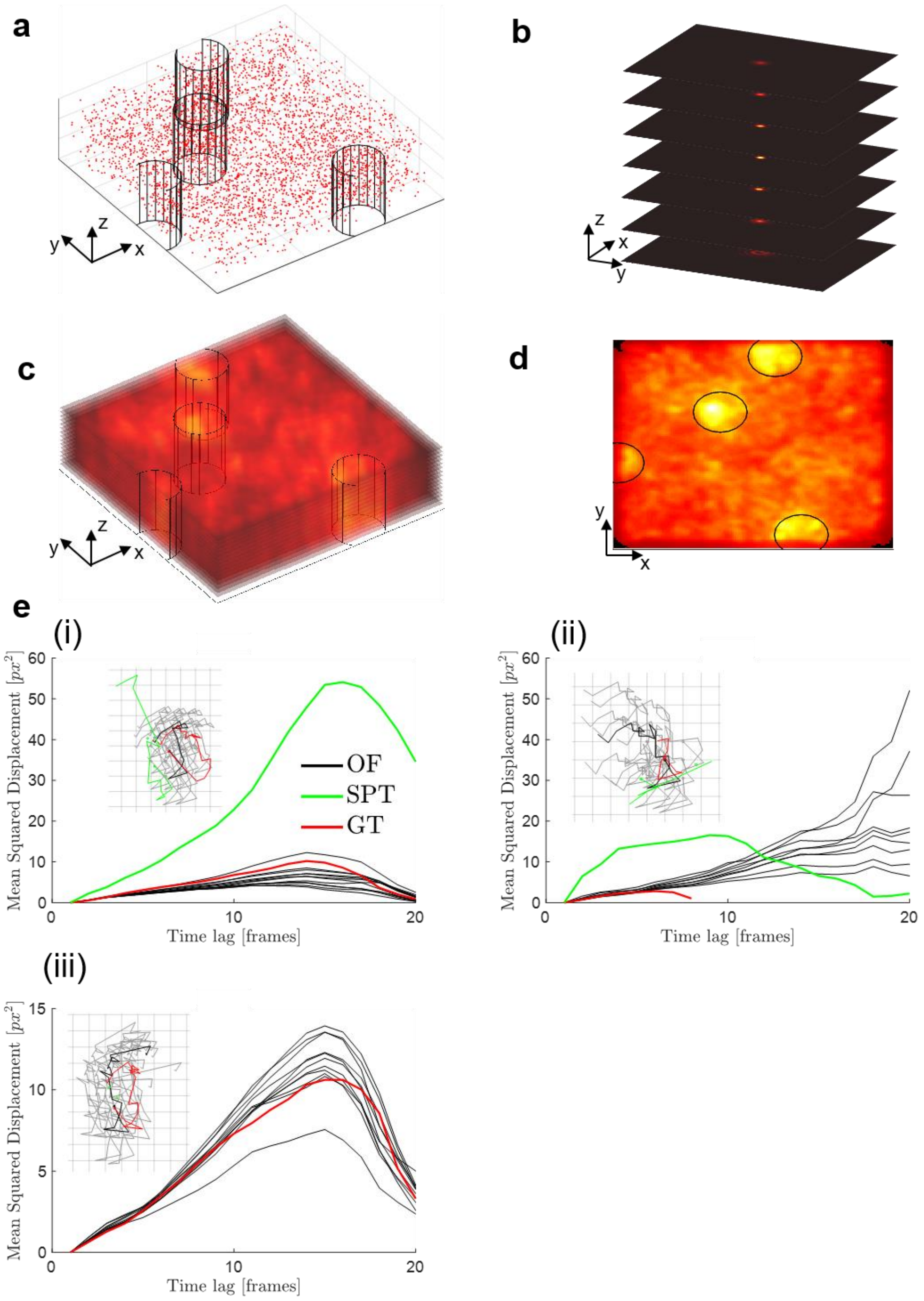


**Fig S5: Simulation workflow and exemplary evaluation. a)** The initial coordinates of 6857 emitters in the first frame of the image series for the scenario of density patches. Regions of super-high density are marked by cylinders. Simulated density is 0.02 and $0.035/px^3$ for regions of high and super-high density respectively. Simulated volume is 128x128x15 pixels. **b)** Simulated point spread function in three dimensions, which is used for convolution of the emitters in a). **c)** Convolved z-stack of the first frame and **d)** 2D projection. **e)** Example MSD curves with corresponding trajectories. The average trajectory reconstructed from OF is shown in black, the single trajectories in light gray. The trajectory origin is indicated by a dot. Visually, agreement is found between the time evolution of trajectories between GT and OF as well as GT and SPT in parts of the trajectories, whereas in other parts, the trajectories deviate and even propagate in opposite directions. However, the MSD curves of GT and OF estimation largely overlap indicating that the magnitude of displacement is accurately estimated. (i) Similarity of the direction of OF and SPT trajectory for approximately the first half of the trajectory, whereas the GT trajectory propagates in opposite direction. For long time the OF trajectory resembles the GT, whereas the SPT trajectory deviates largely, possibly due to a reconnection error. The MSD curve reflects the similarity of GT and OF estimation despite of the deviation in direction of the trajectories' propagation. The reconnection error in SPT leads to a tremendous increase in MSD values especially for long time lags. (ii) Disappearance of the GT emitter after 7 frames. However, both OF and SPT detect trajectories throughout the whole frame series. (iii) A false negative of SPT and therefore no dynamic information for the SPT data set. f) SPT detections for low (i) and high (ii) density of emitters. SPT is able to reliable detect emitters for low density; however, for high density the emitter signals largely overlap such that a detection of single emitters is impossible. However, due to correlated motion of emitters it is nevertheless possible to extract valuable information about emitter dynamics.

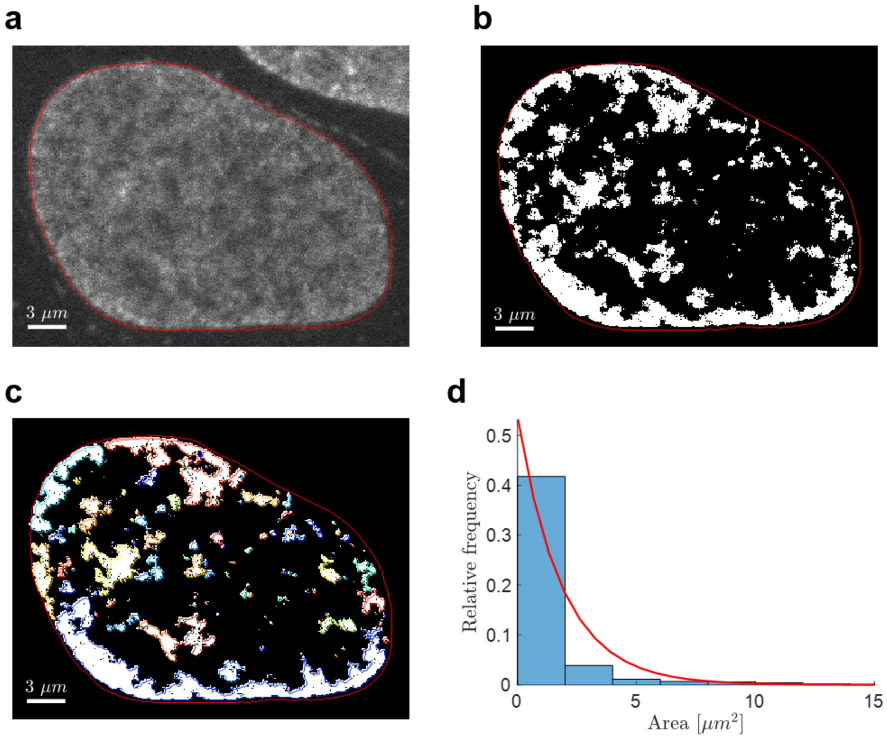

**Fig S6: Size distribution of heterochromatin regions. a)** Raw fluorescence image with nucleus boundary indicated by a red line. **b)** The ~12.5% pixels with highest intensity were found in a Gaussian filtered image as described in [8]. **c)** The independent heterochromatin regions are found (labeled in random color for visualization) and their area is calculated. **d)** The area distribution of heterochromatin domains (blue histogram) can be described by an exponential distribution with mean $(1.9 \pm 3.5)\ \mu m^2$ (red line).

**Performance evaluation**

The simulated image series were analyzed using OF and SPT. The results from 10 independent simulations are summarized in Figure 2a, b of the main text and in Fig S7 for the three scenarios. For a low density of emitters (Fig S7a), MTT is able to detect and reconnect the majority of GT emitters ($79 \pm 4$ % true positives) and yields good performance in all three measures considering value and shape of the estimated MSD curves. In particular, estimated diffusion coefficients show a very low error (< 10%). The OF estimation for relatively sparse signals is difficult leading to comparably lower performance than SPT (relative error in diffusion constant is about 25%). For a high density of emitters (Figure 2a, Fig S7b), SPT performance in terms of the relative Euclidean distance drops, whereas OF performs better than in the low density scenario. However, the relative error in diffusion constant increases for both methods (60% and 40% for SPT and OF respectively). Introducing a density heterogeneity, the SPT performance stays unaffected, but the relative error in diffusion constant for the OF reconstruction is reduced to about 25%. OF estimation uses the bulk motion of many particles such that the vast majority of pixels carries information and the flow field does not have to be approximated by the smoothness assumption underlying OF as in a low density scenario. However, an additional structure in the image such as a density heterogeneity leads to considerably more accurate results than with a uniform density due to additional structure in the image.

These results confirm that the SPT method under consideration is especially accurate when comparably sparse signals are present. SPT is naturally limited to the detection of single emitters or aggregates of emitters [12,13] and therefore lacks accurate information when detection is impossible, e.g. due to high overlap of independent emitter signals. The Optical Flow algorithm in this study is used in high-density scenarios and additional structures in the image such as a heterogeneous density of chromatin enhance the performance considerably. Hi-D therefore constitutes a complementary approach to extract dynamic information of biomolecules with dense labeling where SPT cannot be applied.

It has to be stressed that the simulations carried out incorporate two important aspects of the approach. Diffusion of emitters in and out of focus as well as appearance and disappearance of emitters as well as heterogeneous chromatin structures (i.e. varying chromatin compaction levels) are considered in all simulations. Thus, the values provided are likely to reflect errors associated with experimental data.

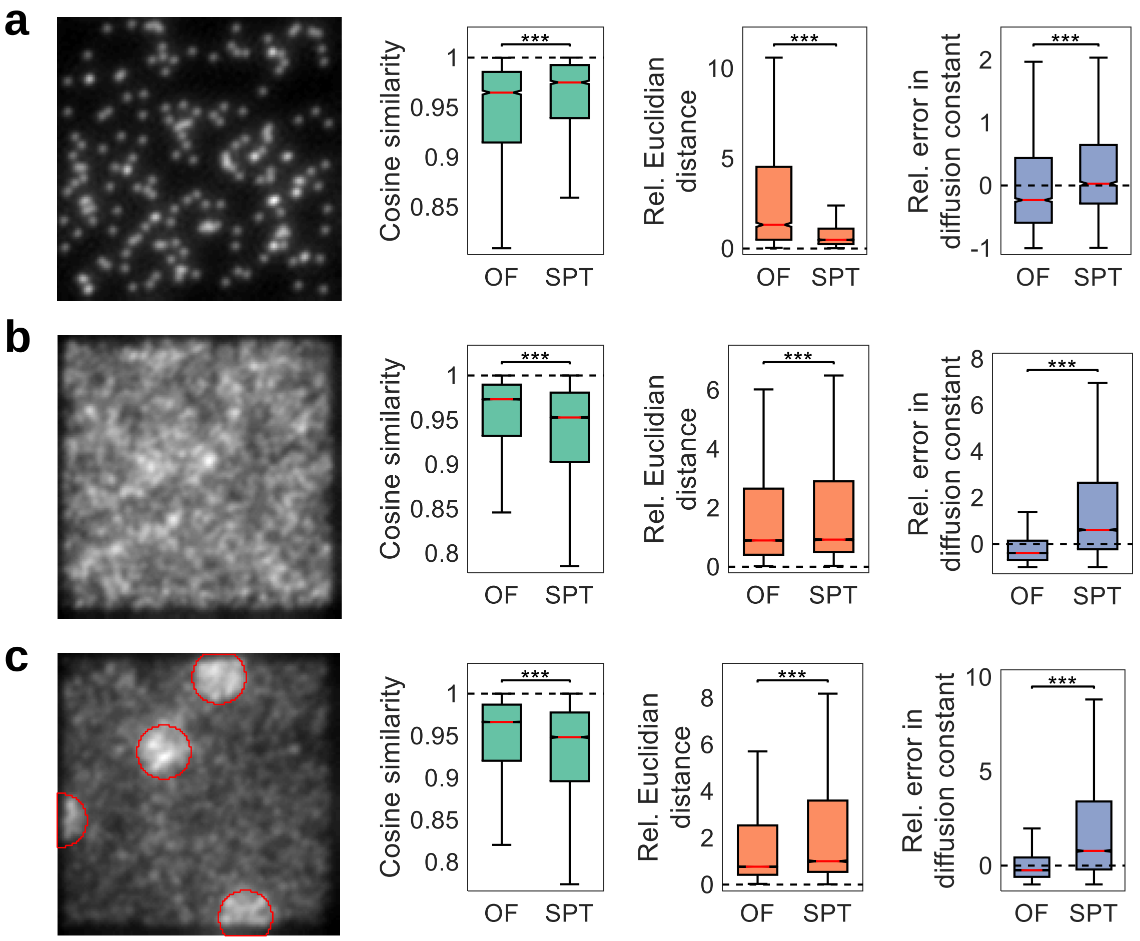


**Fig S7: Comparison of Optical Flow and Single Particle Tracking by simulated image series. a)** First frame of a time series of 20 frames with low density ($0.001/px^3$) of emitters undergoing Brownian motion in 3 dimensions convolved by a typical point spread function. The time series is subject to OF and SPT estimating the trajectories of emitters. From the estimated trajectories, the MSD is computed and compared to the ground truth. Three performance measures are used: Cosine similarity, relative Euclidian distance and the relative error in the determined diffusion constant as determined from fitting. Dashed lines show the optimal value, i.e. perfect agreement between estimation and ground truth. Red lines indicate the median value. Data from 10 independent simulations; outliers are not shown for simplicity. For low density, SPT is able to detect and reconnect emitters over time yielding in more accurate estimates for the MSD compared to OF, which suffers from sparse signal. **b)** High density ($0.02/px^3$) of emitters and the same performance measures as in a). For high density,

OF outperforms SPT in terms of the performance measures. SPT cannot detect particles due to the high overlap of emitters. For the few detected particles, dynamics can be extracted, yielding however overall lower accuracy as the dense motion reconstruction using OF. **c)** High density as in b) with patches of super-high density ($0.035/px^3$) encircled for visualization, imitating regions of densely packed chromatin. For only a small fraction of the image exhibiting a very high density, the superiority of OF in contrast to SPT becomes more evident and even the error in diffusion coefficient determination decreases significantly from b) to c) ($p < 0.001$, data not shown) showing additional strength of OF in local high-density scenarios. Statistical significance assessed by a two-sample Kolmogorov-Smirnov test (*: $p < 0.05$, **: $p < 0,01$, ***: $p < 0.001$). Parts of this figure are reproduced in Figure 2a, b of the main text.

#### NOTE S4

##### On the origins of possible discrepancies between SPT methods and Hi-D

SPT detects spots in each frame and connects them from frame to frame in a certain method-specific way to yield trajectories. Therefore, the loci of interest should be distributed relatively sparse and should be bright (at least much brighter than the background) to ensure reliable spot detection. A bright spot can be easily seen under the microscope even if chromatin 'covers' the spot (that is even if there is chromatin in the focal volume between the labeled locus and the objective). Photons emitted from the labeled locus are not completely absorbed by the surrounding nuclear volume such that the spot can still be detected – note that this is desired in SPT experiments.

In contrast, Hi-D does not aim to detect spots and therefore the complete chromatin is labeled. In an experiment in which a locus is specifically labeled such that a SPT experiment can be carried out and the surround chromatin is labeled simultaneously, one could thus infer the locus position in the image from the SPT spot. However, in the whole-chromatin channel, chromatin above and below (within the focal volume) is equally visible and thus influences the flow field reconstruction. This will inevitably lead to discrepancies between SPT and Hi-D due to the fact that the focal plane is not infinitesimal thin.

To illustrate the issue, a simulated chromosome [14,15] is employed, from which microscopy image series (50 frames covering 10 s) corresponding to our experimental imaging conditions are generated. A random locus (5 kb) is chosen whose photon emission is enhanced 250-fold compared to the surrounding chromatin (Movie S1, left column). A corresponding image series for which the complete genome is homogeneously labeled was generated (Movie S1, middle column). The ground-truth displacement and flow field for the SPT and Hi-D image series, respectively, are superimposed (Movie S1, bottom row). The overlay of SPT displacements and flow field shows that, in general, the displacement vectors at the SPT spot do not agree (Movie S1, right column), thus making the SPT and Hi-D dynamics not directly comparable. Note that the displacements and flow fields shown here are not inferred by SPT or Hi-D, but display the ground truth which is known from the simulation – there is thus no error associated to inaccuracies of either SPT or Hi-D and Movie S1 hence illustrates the inherent difficulty to compare SPT and Hi-D under realistic imaging conditions.

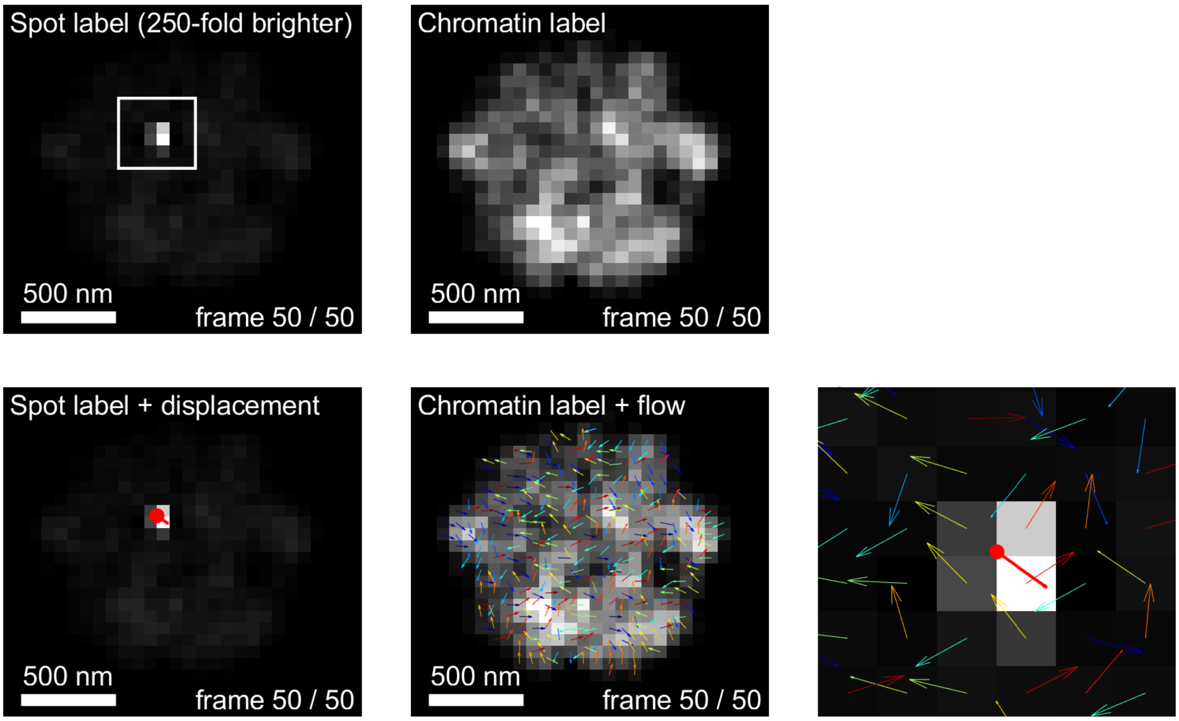


**Movie S1: Illustration of discrepancies due to different imaging modalities between SPT methods and Hi-D using a simulated chromosome.** Fluorescence microscopy images are generated from a simulated chromosome [14,15], corresponding to our experimental imaging conditions. A random locus was simulated 250-fold brighter than the surrounding chromatin in order to mimic a single locus label suitable for SPT analysis (left column). In contrast, a whole chromatin imaging suitable for Hi-D analysis is simulated in the middle column. The lower row depicts the ground-truth locus displacement for the SPT image series and flow field for the Hi-D image series, respectively. A zoom around the single locus spot (right column, depicted by the rectangle in the left column) shows that the single locus displacement vector does not correspond to the local flow field of the genome.

#### NOTE S5

##### Comparison of Hi-D to an orthogonal approach, iMSD

iMSD is usually applied to a region of interest, over which average parameters for the region of interest are extracted. To compare the Hi-D and iMSD approaches, iMSD was applied to successive ROIs spanning an entire nucleus with large overlap, such that parameters were computed for each pixel separately. The resulting maps of diffusion constants showed that regions of high and low mobility were apparent in both methods (Figure 2e, yellow regions marked with red arrows), thus demonstrating qualitative agreement between the two conceptually different methods. iMSD could not compute parameters for ROIs at the nuclear periphery, at which diffusion is expected to be exceptionally low, and in regions of very high mobility (Figure 2e, right, surrounded by yellow-colored pixels). Quantitatively, both methods yield diffusion constants in the same order of magnitude (Hi-D $(1.6 \pm 0.8) \cdot 10^{-3}\ \mu m^2/s$ , iMSD $(2.2 \pm 4.5) \cdot 10^{-3}\ \mu m^2/s$, mean $\pm$ standard deviation). However, we noticed a bias towards very small ($< 5 \cdot 10^{-4}\ \mu m^2/s$), and the presence of a few large ($> 5 \cdot 10^{-3}\ \mu m^2/s$) values, and a considerable spread of the distribution compared to values extracted by Hi-D (Figure 2f). Likewise, values of the anomalous exponent extracted by iMSD showed many spurious values around $\alpha \sim 0$ and $\alpha \sim 2$, while the distribution of values derived from iMSD is reasonable with respect to the magnitude of extracted parameters ($0.25 < \alpha < 1.5$, Figure 2g). We conclude that the estimation of dynamic parameters using Hi-D yields parameters which are qualitatively and quantitatively similar to an orthogonal approach, iMSD, and is thus an accurate read-out of chromatin dynamics. By making use of a Bayesian inference to select a suitable interpretive model for regression of MSD curves, Hi-D is advantageous to existing methods to yield conclusive and spurious-free parameter estimates within the entire nucleus simultaneously. It is to mention that iMSD is applied to ROIs spanning $\sim 50\ \mu m^2$ [16] (about 70 x 70 pixels in our data), while Hi-D uses only a 3 x 3 pixel neighbourhood to estimate the correct diffusion model and its parameters and thus offers greatly enhanced spatial specificity. Finally, we observed a $\sim 35$ speed-up in computational time of Hi-D compared to iMSD for a whole nucleus (roughly 500 $\mu m^2$) with single-pixel resolution.

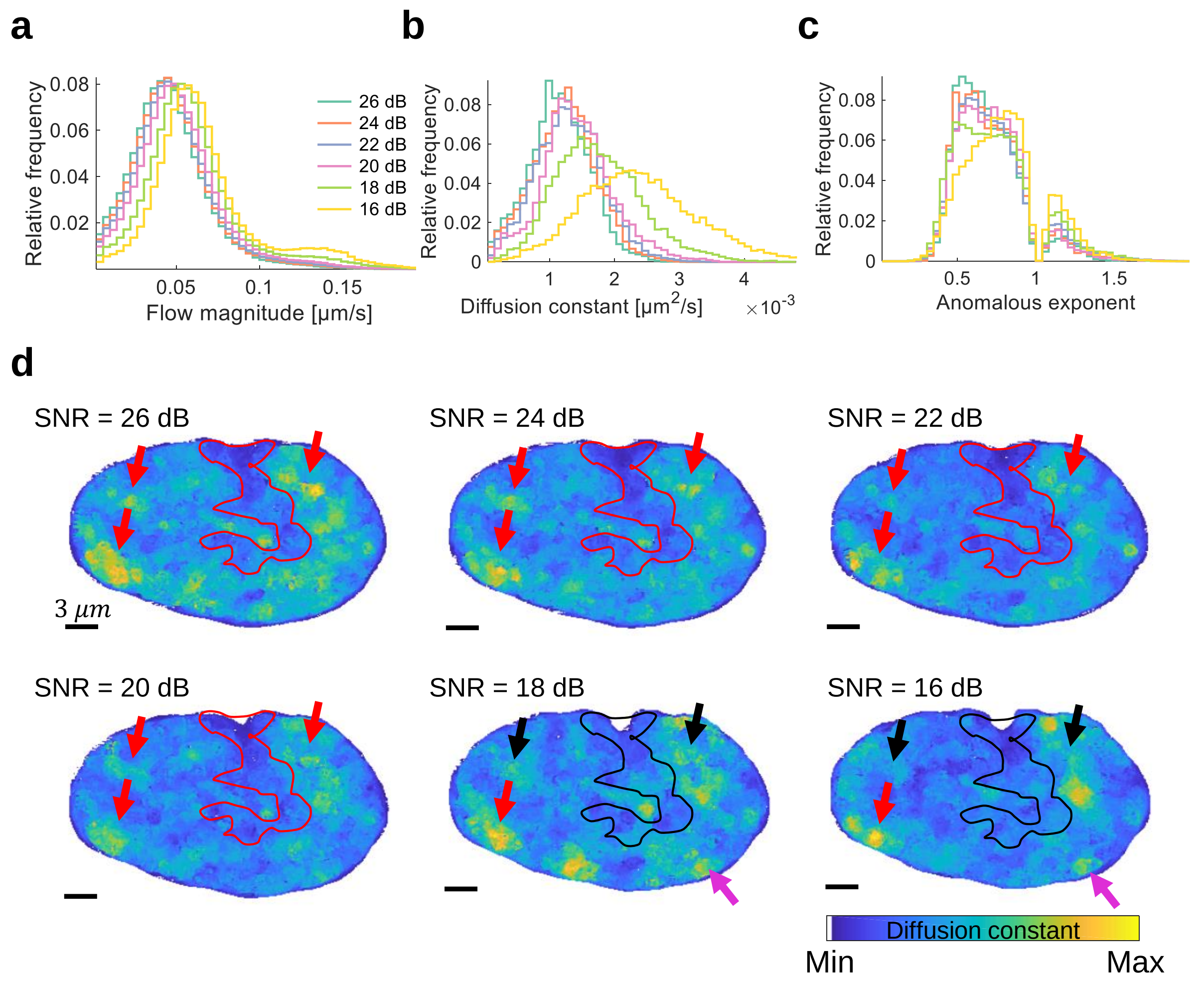


**Fig S8: The effect of varying signal-to-noise ratios on the estimation of dynamic parameters with the Hi-D approach. a)** Flow magnitude, **b)** diffusion constants and **c)** anomalous exponent derived from a nucleus corrupted with varying levels of signal-to-noise ratio (SNR). Results are consistent up to a lower bound of ~ 20 dB. **d)** The spatial distribution of diffusion constants of a representative nucleus is shown after the fluorescence time series of the nucleus was corrupted with varying levels of noise. Features of the diffusion map such as local regions of high mobility (arrows) and traces throughout the nucleus with reduced dynamics (line) are marked. For SNR up to ~20 dB, these features are well conserved. For even lower SNR values, however, some features disappear and/or become faint (black arrows) and originally defined shapes become washed out (black line). Only originally large and/or very mobile regions are conserved for SNR values as low as 16 dB (red arrow in panels with SNR = 18 dB and SNR = 16 dB). Furthermore, noise-related artifactual features arise below 20 dB (purple arrows).

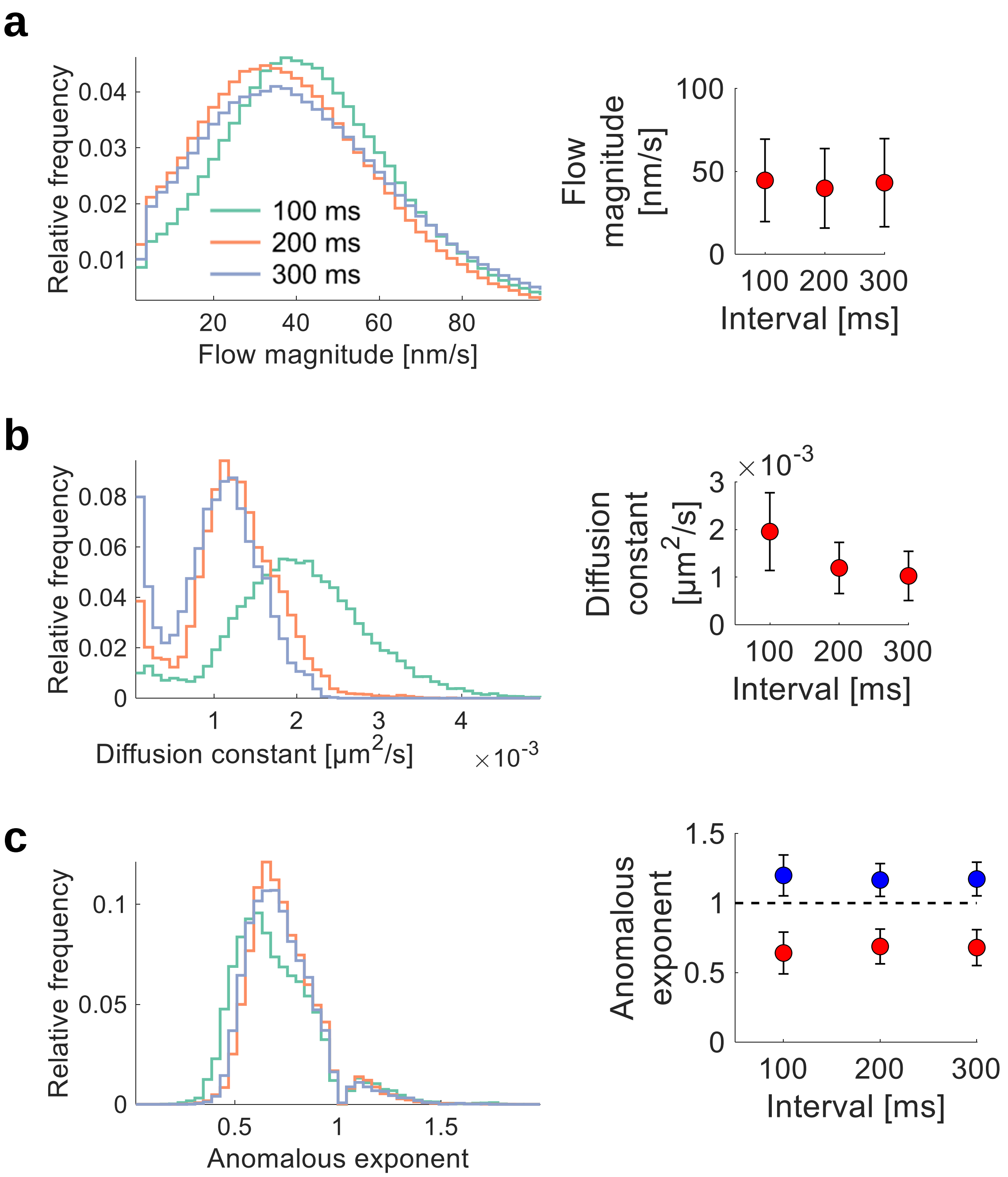


**Fig S9: The effect of varying time intervals between frames on the estimation of dynamic parameters with the Hi-D approach. a)** Flow magnitude, **b)** diffusion constants and **c)** anomalous exponent computed from time series which were images with time intervals of 100 ms, 200 ms and 300 ms between frames, respectively. The right panels show average values and standard deviations from the distributions. The three parameters are consistent when images with 200 ms and 300 ms interval between frames. However, when imaged with 100 ms, distributions deviate: In particular, the diffusion constants are over-estimated (about 2-fold) and the distribution of diffusion constants considerably broadens. This is likely due to the fact that chromatin dynamics become hardly detectable at very short time scales using conventional microscope-setups featuring cameras. As a rough estimate, consider an average diffusion constant of about $\mathrm{D} \approx 4 \cdot 10^{-3}\ \mu m^2/s$ (compare, e.g. Figure 4 of the main text). Displacements between frames are estimated to be in the order of $\sqrt{4D\tau} \approx 40\ nm$, considering simply free diffusion in two dimensions, which is about 60 % of the pixel size (pixel size 65 nm). Detection of dynamics which whose typical length scale between frames is purely in the sub-pixel range of the imaging system is thus difficult to estimate (an exception is camera-based fluctuation correlation spectroscopy, which may estimate local dynamics at shorter length scales), which explains the overestimation of dynamics for acquisition with 100 ms interval between frames. For comparison, a time interval of 200 ms leads to a typical displacement of $\approx 60\ nm$, which is about the dimension of a pixel. An estimation about the expected magnitude of dynamics is thus useful to determine acquisition parameters yielding unbiased results. Note that the considerations above are rough estimated concerning average values and that dynamics can be detected at the sub-pixel level using camera-based imaging systems, if, in average, displacements between frames are about one pixel size. This is possible, because Hi-D uses the collective fluorescence information of the whole nucleus simultaneously to infer local sub-pixel dynamics.

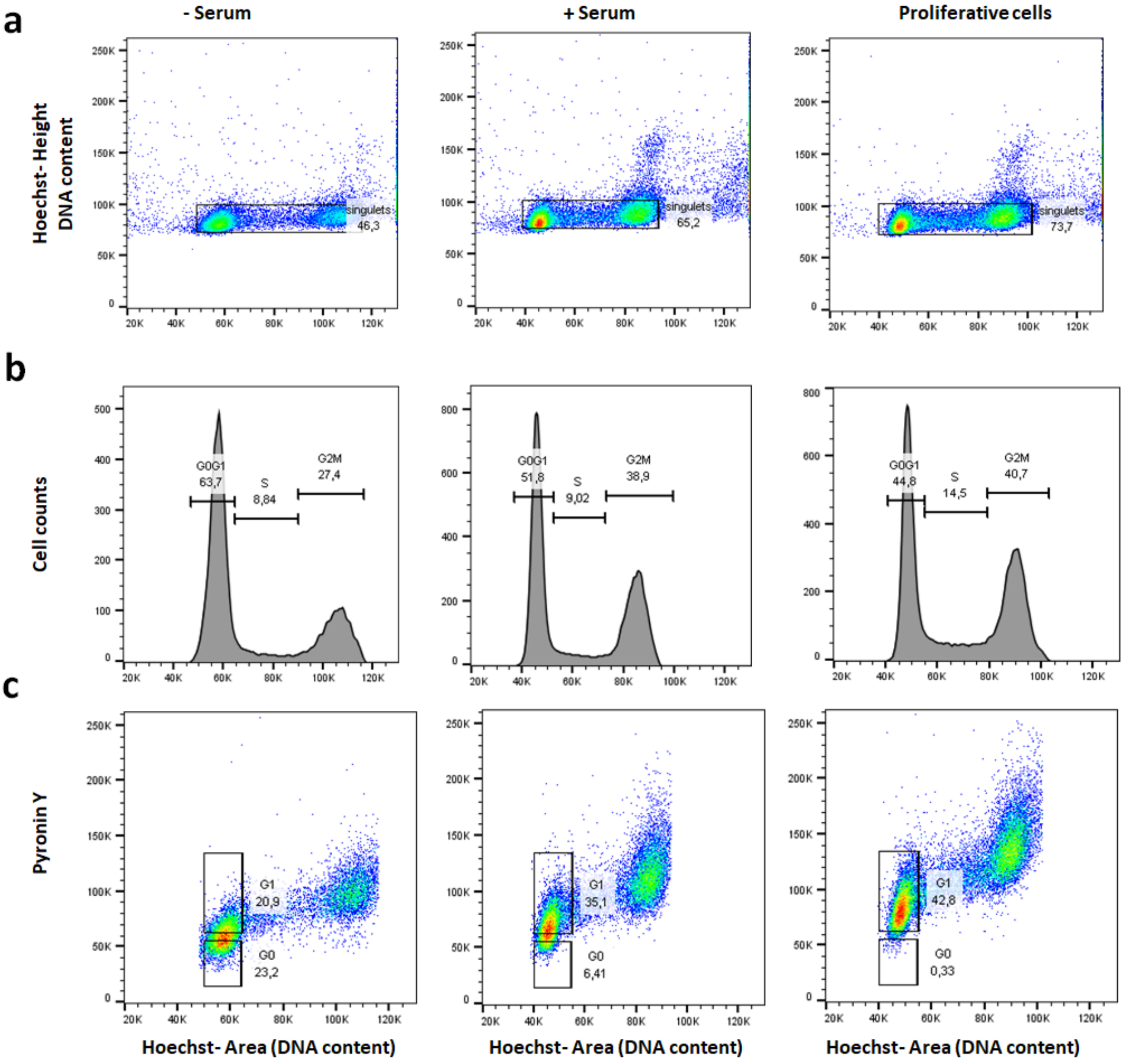


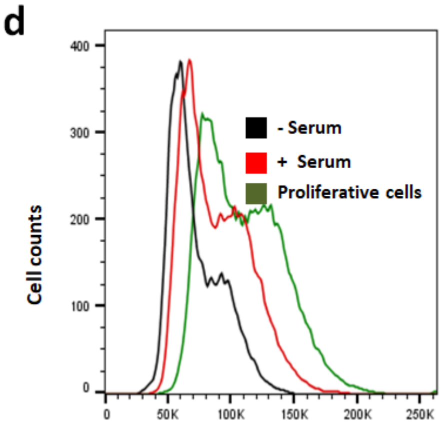


| | Pyronin Y Fluorescence Median |
|---|---|
| **- Serum** | 65.984 |
| **+ Serum** | 84.032 |
| **Proliferative cells** | 105.920 |

**Fig S10: Addition of serum increases transcription without affecting drastically cell cycle phases.** FACS analysis was performed on U2OS cells (<80% confluence) labelled with Hoechst and Pyronin Y to quantify the DNA and mRNA content, respectively. **a)** Singlet events are gated on Hoechst Height vs Hoechst area bivariate in three conditions (serum starved for 24h (- Serum)), addition of serum during 15 min (+ Serum) and normal growth condition (proliferative cells). **b)** DNA content distribution of singlets is used to assign cell cycle phases, in the three conditions mentioned in a). Percentages of cells in cell phases are given. DNA content distributions were performed based on singlet events gated as in a) with Hoechst 33342 labelling. **c)** Singlet events gated by Hoechst 33342 and Pyronin Y fluorescence in the three conditions mentioned in a), as indicated. $G_0$ cells were shown to have less mRNA content compared to G1/S/G2/M cells. For the same DNA count, a low level of Pyronin Y shows $G_0$ cells and a high level of Pyronin indicates G1 cells. $G_0$ and $G_1$ phases are differentiated by empirically drawn boxes. **d)** Distribution of Pyronin Y fluorescence in the three conditions mentioned in a), show an increase in mRNA content (median fluorescence intensity shown in table) in cells after 15 min serum addition. Two independent experiments were performed and were highly reproducible. Results from one replicate are shown.

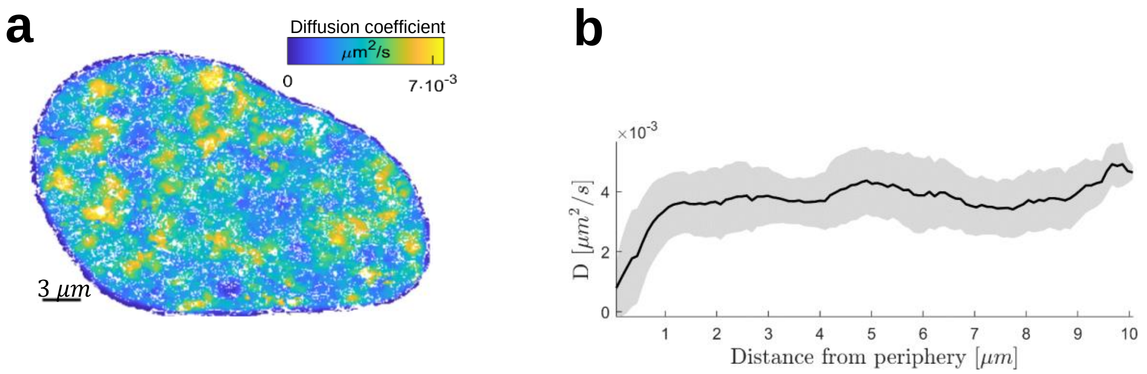


**Fig S11: Chromatin diffusion is reduced at the nuclear periphery compared to the nuclear interior. a)** A representative map of the diffusion constant of chromatin dynamics within a nucleus. **b)** The diffusion coefficient is measured in dependence of the distance to the nearest periphery pixel by a binned Euclidean distance transform (shown is mean value ± standard deviation, n = 13 quiescent cells). The trend holds equally true for actively transcribing cells (data not shown).

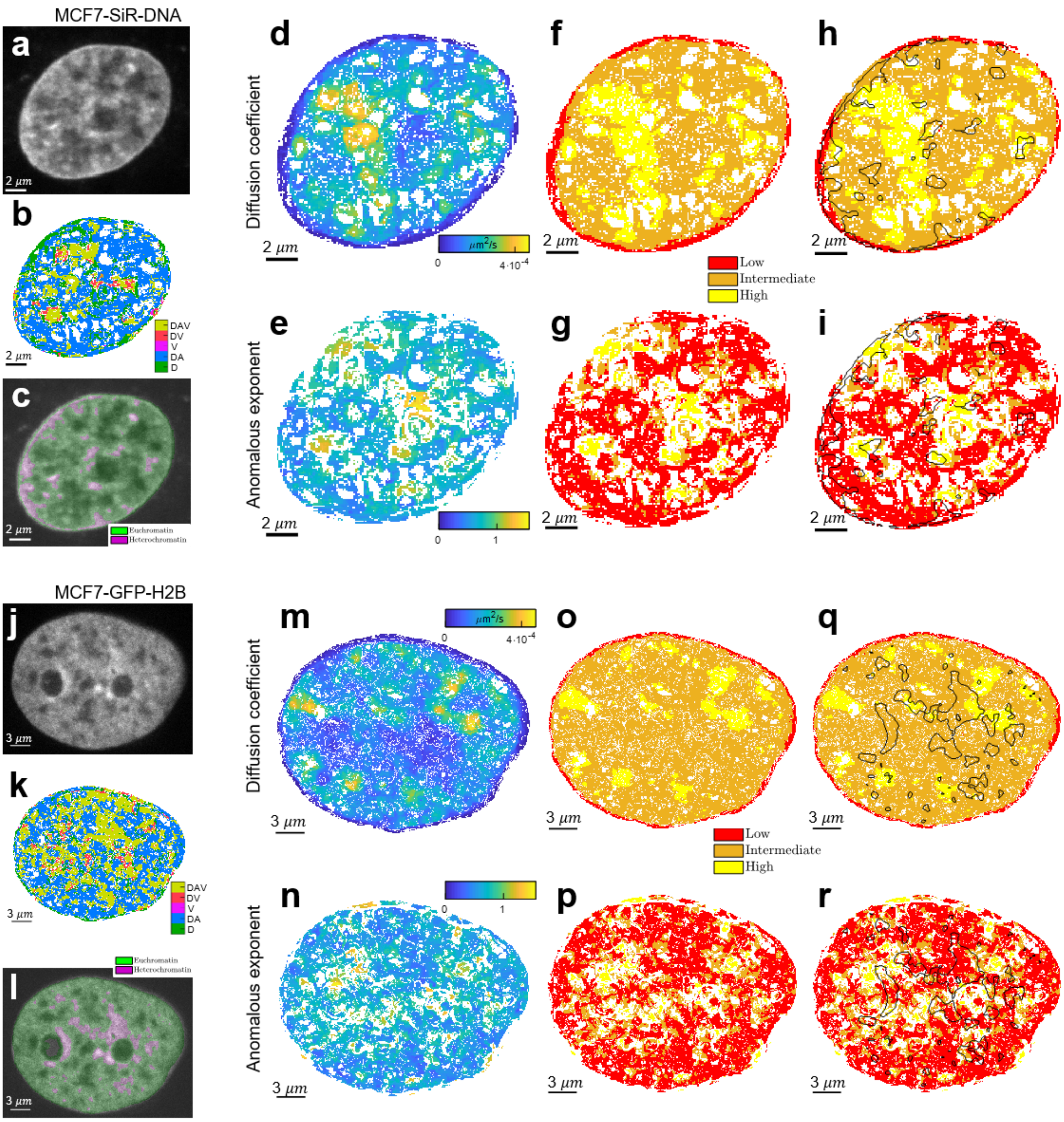


**Fig S12:** Hi-D analysis for a different cell line (MCF-7 cells). **a)** Fluorescence image of a MCF-7 cell nucleus stained by SiR-DNA to which Hi-D was applied. **b)** Model selection as indicated in the color bar. **c)** Spatial discrimination of high- and low compaction indicating hetero- and euchromatin respectively. **d)** The diffusion coefficient and **e)** anomalous exponent extracted from Hi-D. **f-g)** Mobility groups partitioning DNA dynamics within the nucleus for f) diffusion coefficient and g) anomalous exponent, respectively. **h)** Overlay of diffusive populations, **i)** anomalous exponent. **j-r)** As for a-i) for a MCF-7 cell nucleus stained by GFP-H2B.

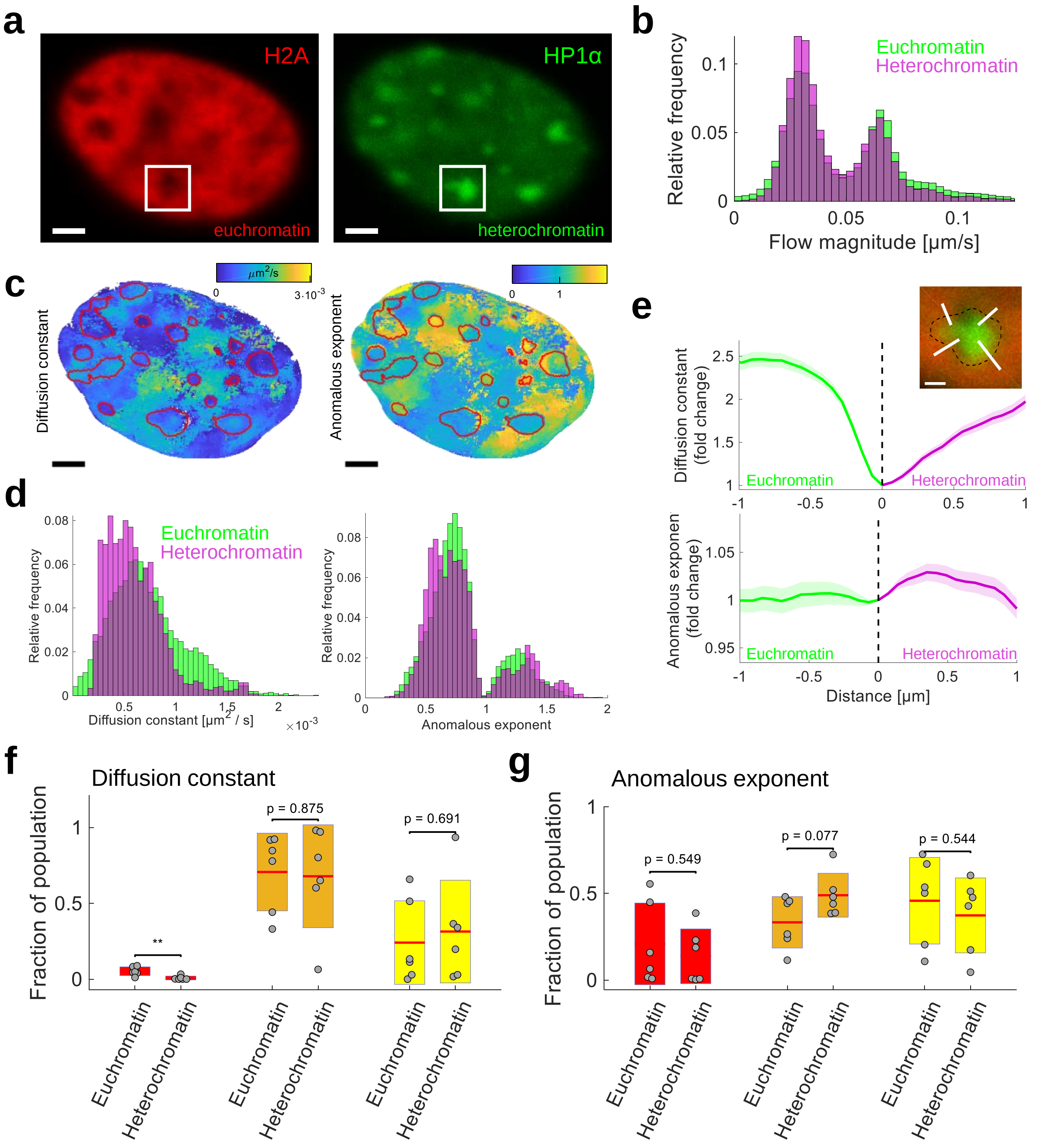


**Fig S13: Dynamics of H2A and HP1α in NIH3T3 cells confirm that chromatin dynamics is independent from compaction, but suggest that eu- heterochromatin phase-separation forms a barrier for chromatin mobility. a)** Exemplary NIH3T3 cells, in which euchromatin was visualized with mCherry-H2A (left) and heterochromatin is visualized by GFP-HP1α (right). Labeling of H2A and HP1α allows to directly assess the presence of heterochromatin. Enhanced HP1α fluorescence and therefore heterochromatin was detected (right) by simple thresholding. Scale bar is 3 µm. **b)** The flow field for a time series of 50 frames was computed and the flow magnitude is shown for eu- and heterochromatin in green and purple, respectively. The distributions coincide indicating that compaction does not directly influence chromatin dynamics from frame to frame. **c)** The diffusion constant (left) and anomalous exponent (right) were computed and are shown exemplary for the cell in A. Heterochromatin is surrounded by red lines. Scale bar is 3 µm. **d)** Histograms of the diffusion constant (left) and anomalous exponent (right) in eu- and heterochromatin. The distributions are qualitatively similar, however, heterochromatin exhibits a narrower range of diffusion constant values than euchromatin. No difference between distributions of the anomalous exponent between eu- and heterochromatin can be observed. **e)** Phase-separation of eu- and heterochromatin, more specifically of HP1α as suggested previously (Strom et al, Nature), would give rise to an observable boundary across which diffusion is hindered. Inside the phases (eu- and heterochromatin), however, chromatin dynamics must not differ necessarily. To prove the existence of such a boundary using Hi-D, the diffusion constant (upper panel) and anomalous exponent (lower panel) were drawn at lines perpendicular to heterochromatin-defining boundaries (inset, overlay of H2A and HP1α of the rectangle indicated in a). A boundary at which the diffusion constant drops about 2.5-fold compared to bulk euchromatin is observed and chromatin dynamics is recovered about 1 µm from the boundary in the interior of phase-separated heterochromatin droplets. While the diffusion constant is strongly reduced, no effect of the boundaries on the anomalous exponent was observed. Scale bar is 1 µm. **f)** A tendency of a mobility or **g)** anomalous exponent population to be associated with either eu- or heterochromatin could not be observed. Statistical significance assessed by a two-sample t-test (*: $p < 0.05$, **: $p < 0.01$, ***: $p < 0.001$).

#### SUPPLEMENTARY REFERENCES

1. Shaban HA, Barth R, Bystricky K. Formation of correlated chromatin domains at nanoscale dynamic resolution during transcription. Nucleic Acids Res. 2018;46.

2. Donea J, Huerta A, Ponthot J, Rodr A. Arbitrary Lagrangian – Eulerian Methods. Encycl Comput Mech. 1999;1–25.

3. Qiu G, Henke S, Grabe J. Application of a Coupled Eulerian-Lagrangian approach on geomechanical problems involving large deformations. Comput Geotech. 2011;38:30–9.

4. Onu K, Huhn F, Haller G. LCS Tool: A computational platform for Lagrangian coherent structures. J Comput Sci. 2015;7:26–36.

5. Jin S, Haggie PM, Verkman AS. Single-particle tracking of membrane protein diffusion in a potential: Simulation, detection, and application to confined diffusion of CFTR Cl - channels. Biophys J. 2007;93:1079–88.

6. Saxton MJ, Jacobson K. Single-particle tracking: applications to membrane dynamics. Annu Rev Biophys Biomol Struct. 1997;26:373–99.

7. Chenouard N, Smal I, de Chaumont F, Maška M, Sbalzarini IF, Gong Y, et al. Objective comparison of particle tracking methods. Nat Methods [Internet]. 2014;11:281–9. Available from: http://www.pubmedcentral.nih.gov/articlerender.fcgi?artid=4131736&tool=pmcentrez&rendertype=abstract%5Cnhttp://www.nature.com/doifinder/10.1038/nmeth.2808

8. Wachsmuth M, Knoch TA, Rippe K. Dynamic properties of independent chromatin domains measured by correlation spectroscopy in living cells. Epigenetics and Chromatin. BioMed Central; 2016;9:1–20.

9. Zidovska A, Weitz D a, Mitchison TJ. Micron-scale coherence in interphase chromatin dynamics. Proc Natl Acad Sci U S A [Internet]. 2013;110:15555–60. Available from: http://www.pnas.org/content/110/39/15555.long

10. Shinkai S, Nozaki T, Maeshima K, Togashi Y. Dynamic Nucleosome Movement Provides Structural Information of Topological Chromatin Domains in Living Human Cells. PLoS Comput Biol [Internet]. 2016;12:1–16. Available from: http://dx.doi.org/10.1371/journal.pcbi.1005136

11. HANSER BM, GUSTAFSSON MGL, AGARD DA, SEDAT JW, Francisco CS, Francisco S. Phase-retrieved pupil functions in wide-field fluorescence. J Microsc [Internet]. Wiley/Blackwell (10.1111); 2004 [cited 2018 May 30];216:32–48. Available from:

http://doi.wiley.com/10.1111/j.0022-2720.2004.01393.x

12. Vig DK, Hamby AE, Wolgemuth CW. On the Quantification of Cellular Velocity Fields. Biophys. J. 2016. p. 1469–75.

13. Jaqaman K, Loerke D, Mettlen M, Kuwata H, Grinstein S, Schmid SL, et al. Robust single-particle tracking in live-cell time-lapse sequences. Nat Methods [Internet]. 2008;5:695–702. Available from: http://www.nature.com/doifinder/10.1038/nmeth.1237

14. Qi Y, Zhang B. Predicting three-dimensional genome organization with chromatin states. Ma J, editor. PLOS Comput Biol [Internet]. 2019;15:e1007024. Available from: http://dx.plos.org/10.1371/journal.pcbi.1007024

15. Barth R, Bystricky K, Shaban HA. Coupling chromatin structure and dynamics by live super-resolution imaging. bioRxiv. Cold Spring Harbor Laboratory; 2019;777482.

16. Di Rienzo C, Gratton E, Beltram F, Cardarelli F. Fast spatiotemporal correlation spectroscopy to determine protein lateral diffusion laws in live cell membranes. Proc Natl Acad Sci. 2013;
